# Colloidal DNA nanoaggregates applied towards file-partitioning for information storage and dynamic data obfuscation

**DOI:** 10.64898/2026.08.25.747150

**Authors:** Sneha Mukherjee, Kevin N. Lin, Kevin Volkel, James M. Tuck, Albert J. Keung, Orlin D. Velev

## Abstract

The molecular programmability of nucleic acids has facilitated the development of architected DNA/RNA nanostructures and their applications in novel materials and technologies. We report how different types of DNA and RNA nanoaggregates, bundling digital information encoded into oligo libraries, can be formed by manipulating the ionic strength of the solution. As DNA or RNA suspensions are immersed in solutions of increasing salt concentrations, we observe the onset of aggregation. Further increase in ionic strength leads to the formation of stable, reproducible, and well-defined aggregates. We show that these nanoaggregates are kinetically trapped at room temperature, stably partition DNA libraries that encode image files, and support file-specific random access by bundling DNA libraries with unique address oligos. The nanoaggregate files can be disrupted and reformed into scrambled bundles using simple external fluid shear or temperature annealing, rapidly obfuscating the data. We term these nanoaggregates nucleic acid PACKeTs: Partitioned Aggregates of Colloidal DNA/RNA through Kinetic Trapping. Overall, the results demonstrate how gaining fundamental insights into ionic colloidal aggregation enables new forms of manipulation of DNA and RNA libraries. This understanding could lead to novel functionalities including kinetically trapped data partitioning, random access, and data encryption or obfuscation.

## Introduction

DNA could provide transformative solutions to handle the exponential growth of digital information^1^. The high theoretical information density of DNA and its ability to be replicated and preserved have motivated the development of numerous prototype DNA-based information storage systems^2–6,7^. There are diverse approaches for storage, retrieval, and even computation in DNA; yet, they share a common and foundational reliance on programmable DNA-DNA interactions. However, as systems necessarily scale in size and sequence diversity (upward of 10^15^ distinct strands per cubic centimeter)^8^ the field will need to expand our understanding beyond DNA-DNA hybridization. It will be essential for predicting how complex DNA systems operate at scale and could present new tools and phenomena for engineering useful functionalities.

Beyond simple base-pairing rules, there are two key factors influencing DNA behavior: the ionic strength of the solution and local mechanical perturbations. The influence of counterions and mechanical forces^9–11^ on DNA hybridization have been extensively studied theoretically^12–16^, computationally^17–19^ and experimentally^20–22^. These studies have been leveraged to control the assembly, size, shape, and function of DNA-gold nanoparticles^19,23^, DNA-protein complexes^24,25^, DNA nanocrystal conjugates^22,26^ and DNA origami structures^27–32^. However, ionic strength and mechanical perturbations could also influence DNA behavior beyond hybridization based on base-pairing. For example, other polymers and small molecules are known to form sequence-independent colloidal structures of specific size and morphology by controlling salinity, microflows and other microenvironmental conditions^33^. Supramolecular assembly of DNA by simple buffer manipulations would be of particular interest for DNA storage and computation, as it would be an important ‘side-effect’ to consider but also present opportunities for engineering new functionalities. Observing sequence-independent supramolecular DNA assembly would also have a broader impact on our understanding of native biological processes and commonly used biochemistry techniques such as chromatin immunoprecipitation.

The central theme of this study is to understand the physical properties and sequence-independent molecular aggregation of DNA and RNA in solutions, and to leverage this understanding to imbue new functionalities for nucleic acid storage systems (**Fig. 1**). We discover that the ionic strength of a solution can induce tunable formation of nanoaggregates from single- and double-stranded DNA, as well as RNA. This aggregation occurs independent of their sequences and is stable over long periods of time. However, these nanoaggregates can be disrupted and reformed by applying transient external shear forces or heat. We term these nanoaggregates nucleic acid PACKeTs: Partitioned Aggregates of Colloidal DNA/RNA through Kinetic Trapping. We harness this phenomenon to create kinetically trapped PACKeTs with DNA encoding distinct photos. When two sets of PACKeTs encoding two distinct photos are mixed into the same solution, DNA strands do not interchange between PACKeTs. Including file address oligos within each PACKeT enables each encoded photo to be distinctly recovered from the mixture. However, intense mechanical disruption such as vortexing or heating rapidly disrupts the PACKeTs, and they reform with DNA encoding the two photos now scrambled together, providing a simple and rapid mechanism to physically obfuscate data. The fundamental insights gained from studying colloidal interactions in this research may enable the design of self-assembled, salt- and mechano- actuated DNA/RNA-based nanoarchitectures in abiotic and biotic applications.

**Fig. 1.**
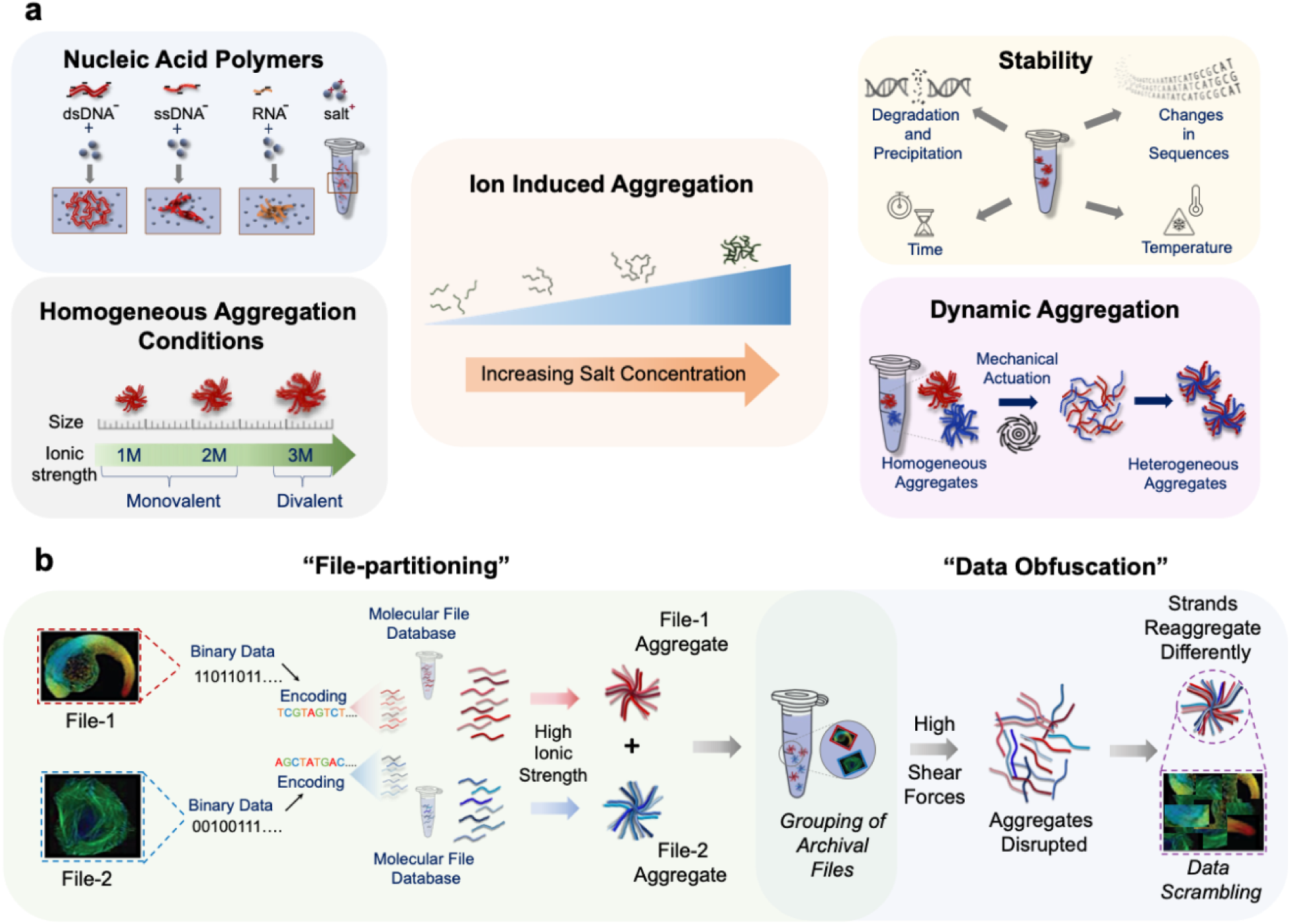
Synthesis and characterization of colloidal nucleic acid nanoaggregates and their application towards information storage. **(a)** Colloidal nanoaggregates of DNA molecules are formed as a result of ion-induced aggregation. Well-defined aggregates of ssDNA, dsDNA and RNA are observed at high concentrations of monovalent and divalent ionic salts. These aggregates resist degradation or disintegration despite variations in external and internal environmental conditions. However, the aggregation is dynamic under certain conditions and can be reversed. (**b)** These salt-actuated aggregates enable two distinct DNA-based data storage functions: file partitioning and data obfuscation.

## Results and Discussion

### Ionic strength drives DNA and RNA nanoaggregation

Salts are ubiquitous in native and synthetic biological systems and, through charge-shielding and other putative mechanisms, are known to affect DNA-DNA hybridization driven by sequence complementarity. We hypothesize that increased ionic strength might also drive non-specific and sequence-independent aggregation of DNA through backbone, sugar, and/or base interactions. Single stranded DNA (ssDNA) differ from double stranded DNA (dsDNA) in the availability of unbound bases and their net charge; we therefore investigate both 120 nucleotide (nt) ssDNA and 120 base pair (bp) dsDNA (**Supplementary Table 1**). Using dynamic light scattering (DLS), we first measure as a baseline the sizes of the DNA dispersed in deionized water (**Supplementary Fig. 1a and b**). The median sizes are ∼10 nm and ∼30 nm for the 120 nt ssDNA and 120 bp dsDNA, respectively. These sizes are consistent with their respective predicted radii of gyration of ∼10 nm and ∼40 nm, based upon persistence lengths of 10 nm and 50 nm and a worm-like chain model for semi-flexible polymers.

Next, DLS measurements of the DNA sizes show that they grow larger with increasing ionic strength, suggesting aggregation is occurring (**Fig. 2a and Supplementary Figure 1c-d**). TEM micrographs verify that aggregation is occurring (**Fig. 2b-c**). Furthermore, the aggregates remain stable at 4°C for at least 7 days without any further growth in size or precipitation suspensions were also stored at 4°C and measured after 1 day and 7 days under the same conditions. The aggregates remained stable for at least 7 days without any phase separation, notable clustering or precipitation (**Fig. 2b-c**). The sizes of the aggregates are reproducible across multiple experiments (**Supplementary Fig. 2**).

**Fig. 2.**
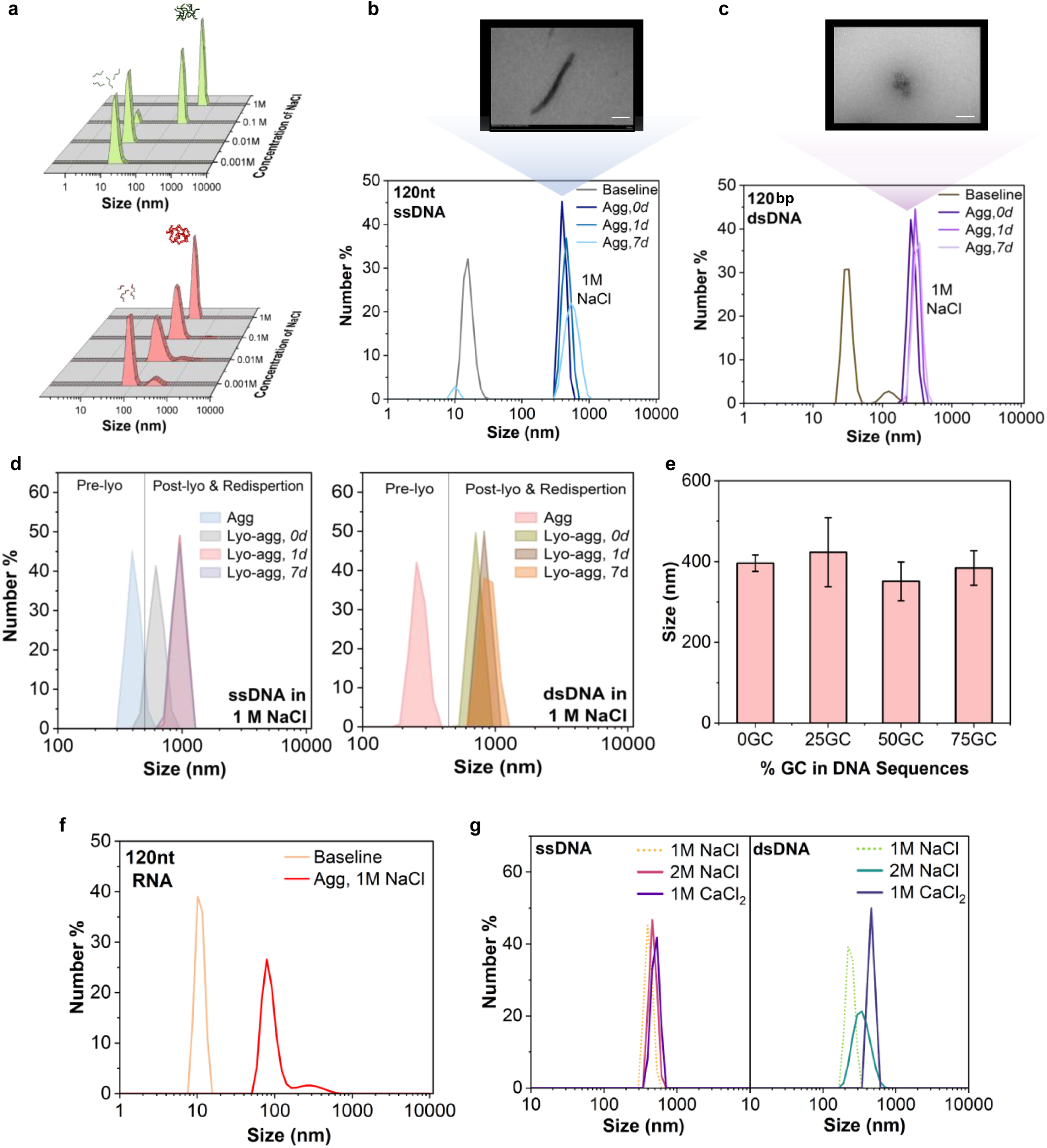
Ionic strength drives DNA and RNA nanoaggregation. **(a)** Z-axis represents number % with peaks indicating the formation of DNA nanoaggregates measured by DLS of **(top)** ssDNA and **(bottom)** dsDNA strands as function of salt concentration. The uniform aggregates of **(b)** 120 nt ssDNA, and **(c)** 120 bp dsDNA, in 1M NaCl remain stable after 7 days. **(inset)** TEM micrographs of the respective aggregates. Scale bars are 200 nm. **(d)** Plots show the comparison of ssDNA and dsDNA aggregates before and after lyophilization. **(e)** Comparison of aggregate sizes for ssDNA samples with sequences of varying GC composition. **(f)** Plot comparing the size of 120 nt RNA in 0 and 1 M NaCl. **(g)** Plots show the effect of increasing ionic strength and divalent ions on ssDNA and dsDNA.

We next investigate how nucleic acid characteristics affect this phenomenon and how robust it is to perturbations. We find that longer strands (200 nt) consistently aggregate in higher ionic strength solutions (**Supplementary Fig. 3**). Furthermore, mixed 120 and 200 nt strands also aggregate to form a unimodal population (**Supplementary Fig. 4**). In addition, DNA sequence does not appear to affect the aggregation behavior. In particular, %GC content can affect DNA flexibility and hairpin formation which may affect aggregation^34^; however, we find changing the %GC content from 0 to 75% does not affect aggregation (**Figure 2e**). We also observe that salt-induced aggregation occurs in RNA systems (**Figure 2f and Supplementary Fig. 5**). These results suggest salt-induced aggregation is a general feature of nucleic acid polymers.

While the DNA aggregates are stable for at least 7 days at 4°C, DNA samples for long-term storage would typically be lyophilized^35–38^. We therefore poise the question if lyophilization would change aggregation behavior. We lyophilize ssDNA and dsDNA and store the samples at - 20°C for up to 7 days and then resolubilize the samples in deionized water. Aggregation remains after resolubilization (**Fig. 2d**). Interestingly, the aggregates of the resolubilized samples are larger compared to samples that were not lyophilized. We do not observe any degradation or precipitation.

It can be expected that the ionic strength generally, and not just NaCl specifically, drives the aggregation. Indeed, we find that both ssDNA and dsDNA aggregate in CaCl_2_ solutions (**Fig. 2g and Supplementary Fig. 6**). Their aggregate size appears to increase with the ionic strength (**Fig. 5g**). With a 3× increase in ionic strength, the average size of assemblies increases ∼1.4× for ssDNA and ∼1.8× for dsDNA. The data for aggregates from ss and ds-DNA of strand length 200 are compared in **Supplementary Fig. 6c**. The size distribution indicates that the addition of divalent cations to DNA solutions leads to strong attraction between the DNA strands, resulting in the formation of very condensed structures.

### Modulating the formation of DNA nanostructures through ionic self-association

The discrete structure of the nanoclusters was investigated by Bio-Transmission Electron Microscopy (Bio-TEM). TEM images for DNA suspended in pure DI, serving as controls, are presented in **Supplementary Fig. 7**. The suspensions of ssDNA and dsDNA in ionic medium revealed distinct nanoaggregate morphologies. We observed that the 120 nt ssDNA aggregate sample adopted configurations resembling “bundles” or “rods”. Given the scale, we refer to these structures as “nanobundles”. These distinct structures are illustrated in **Fig. 4a** and **b** and were consistently observed throughout the grid (**Supplementary Fig. 8**). **Fig. 4c** provides a visualization of the modes of DNA arrangement into these thick bundles. The histogram in **Fig. 4d** represents the size distribution of these bundles measured along their longest dimension, approximated with a normal distribution. The sizes measured by processing the images are in good agreement with the results obtained through light scattering.

The structure and morphology of the dsDNA aggregates under the same conditions was very different. The dsDNA samples formed closely packed globular clusters illustrated in **Fig. 4e and f**. Similarly to the nanobundles, these nanoclusters appeared uniform, stable, and spread throughout the grid. The possible structure of these clusters is visualized in **Fig. 4g**, while **Fig. 4h** represents the size distribution of these clusters approximated with a normal distribution. Additional TEM images are presented in **Supplementary Fig. 8**.

### Ionic Influence on the Dynamic Reassembly of DNA nanoaggregates

We also assessed the reversibility of the aggregation process, which can be important for applications such as information storage and readout. This was achieved by adding two dsDNA (120 and 200bp) samples individually to two systems with higher concentrations of salt: one with 2 M NaCl and the other with 1 M CaCl_2_. The study included three phases as illustrated in **Fig. 3a** and **b**. Initially, we measured the DNA aggregate sizes in both systems. Next, we briefly vortexed the samples to induce mechanical shear and immediately measured the average sizes to assess the impact of shear forces on the aggregates. Finally, the samples were allowed to stand undisturbed for 30 mins, after which measurements were repeated to follow the aggregate size distribution over time.

**Fig. 3.**
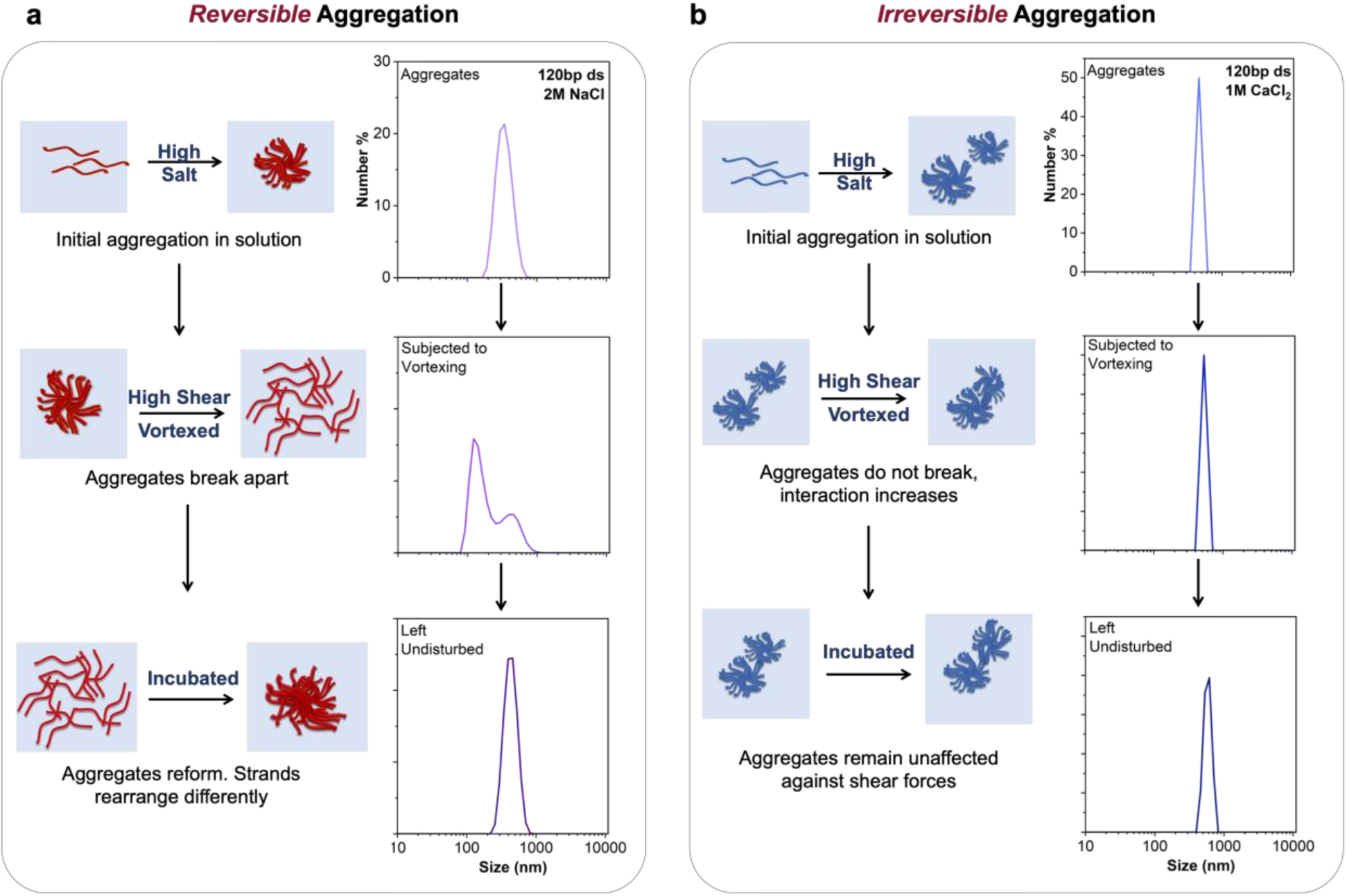
Reversible and irreversible DNA nanoaggregation in well-defined structures. 120 bp dsDNA are subjected to 2 M NaCl and 1 M CaCl_2_. **(a)** This section illustrates the *reversible* aggregation. Schematic shows that the aggregates formed at 2 M NaCl break off on vortexing. When left undisturbed, they reaggregate back to form a similar assembly as the initial one. This is the case of monovalent ions and relatively lower ionic strength (I = 2 M). Plots shows how the DNA aggregates break down and reassemble in 2 M NaCl. Although these reformed aggregates lie within the same size range as the initial aggregate, they can have unique features and characteristics. **(b)** Schematic shows how aggregates at high ionic solutions (I = 3 M) or with divalent ions (Ca^2+^) remain stable and unperturbed, exhibiting an *irreversible* aggregation. DLS data show how these aggregates remain undisturbed and extremely stable with 1 M CaCl_2_.

**Fig. 4.**
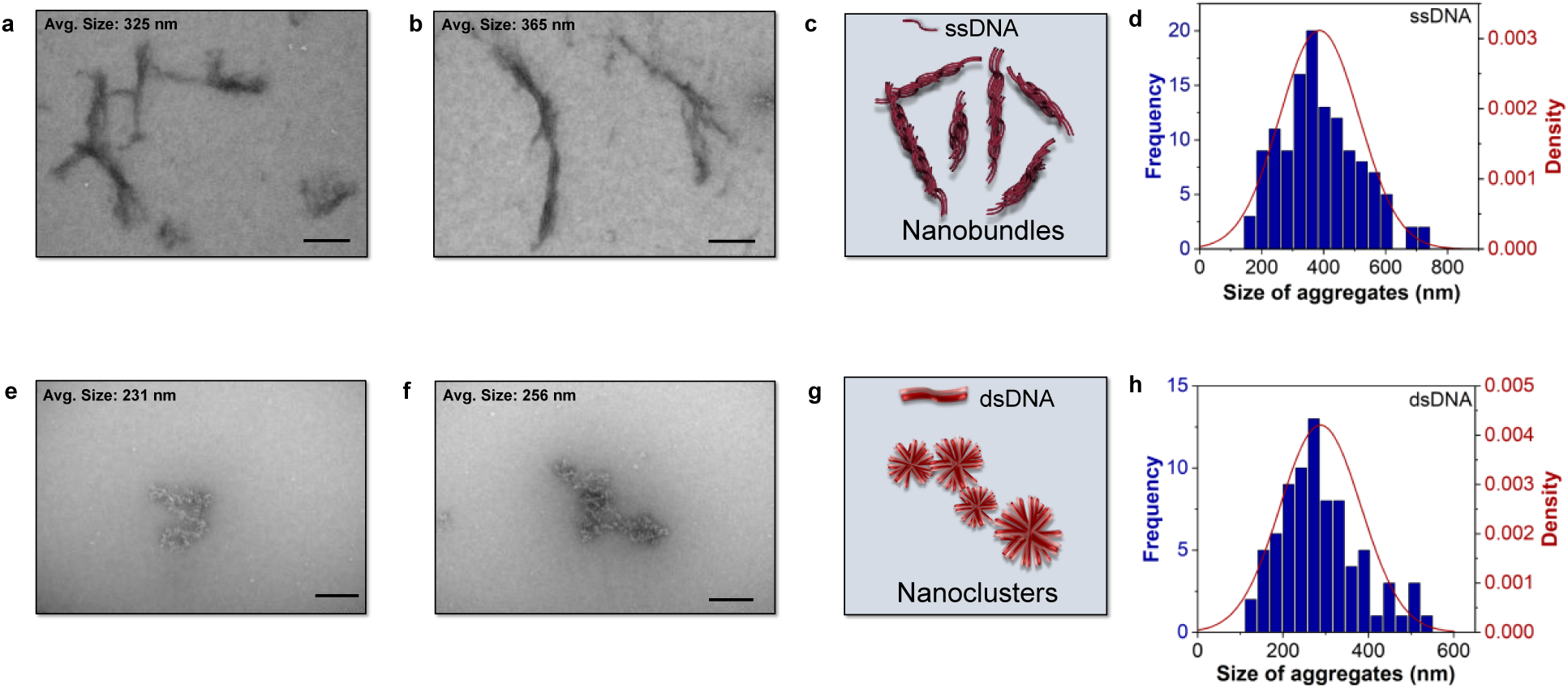
Formation of distinct DNA nanostructures driven by ionic interactions. **(a), (b)** Bio-TEM indicates the formation of aggregates of ssDNA in 1 M NaCl. **(c)** Schematics of the possible arrangements of ssDNA strands. The assembly is referred to as “nanobundles”. **(d)** Size distribution of the nanobundles approximated by a normal curve. **(e), (f)** Bio-TEM indicates the formation of aggregates of dsDNA in 1M NaCl. The images show the “*nanoclusters*” and a possible network of nanoclusters. **(g)** Schematic illustrating the possible arrangement of dsDNA strands. The assembly is referred to as “nanoclusters”. **(h)** Size distribution approximation of the nanoclusters approximated by a normal curve. All scale bars are 200 nm.

Two distinct outcomes were observed. The system of dsDNAs in 2M NaCl exhibited a decrease in the size of the aggregates after vortexing (**Fig 3a**). The peak shifted to smaller sizes and was not as distinct and narrow compared to the initial aggregates. This shift towards the smaller sizes indicated that the aggregates may be breaking apart or getting disrupted. However, when we left the samples undisturbed for 30 mins and measured again, the peak shifted towards a higher size suggesting a process of aggregates re-assembly. However, it is possible that the structure of the re-assembled aggregates is different than the one formed in the first step. In contrast, for dsDNAs in 1M CaCl_2_, there was little to no change in the aggregate behavior and the peak remained unchanged after vortexing (**Fig 3b**). In the case of 2M NaCl, the aggregation is reversible, whereas, in the case of 1M CaCl_2_, the aggregation is irreversible. Other cases with ssDNA are presented in the **Supplementary Fig. 9**.

In addition to using fluid shear to trigger disassembly, we used thermal treatment to disrupt the aggregates. The temperature of each ssDNA and dsDNA sample (120 bp) was increased stepwise from 25°C to 45°C, to 60°C, and finally to 80°C in a continuous process (**Supplementary Fig. 10**). The temperatures were maintained below 80°C to avoid the DNA denaturation. Initially, the aggregates remained in the same size range when we raised the temperature from ambient conditions to 45°C. For both ssDNA and dsDNA, we observed a drop in aggregate size at 60°C, indicating that the aggregates were disrupted at higher temperatures. For dsDNA, we could reverse the aggregation and re-form the aggregates by first heating the sample and then cooling it. **Supplementary Fig. 10** elucidates the temperature-responsive aggregation dynamics of dsDNA aggregates.

### Formation of DNA nanoaggregates can facilitate strand exchange in an ionic condition

Having established the formation of stable DNA nanoaggregates, we assessed the mechanisms of their aggregation-disruption-reaggregation cycle on a molecular scale and explored how these DNA nanoaggregates could enhance DNA-based data storage, leveraging their on-demand assembly and reassembly. The first step was to check the stability and efficacy of the aggregates during pull-out tests using single oligos. We used two 200 nucleotide (nt) single-stranded DNA oligos (ssDNA, ssS1 and ssS2) with distinct primer binding sequences at the 3’end. Both oligos also contained a common 23 nt sequence, which was inserted 20 nt from the 3’end. We used this common sequence to convert the ssDNA into double-stranded DNA with an overhang extension (ovDNA, ovS1 and ovS2), following the protocol described in a previous system^6^. The primer binding sequence allowed for the hybridization of a biotin-labeled primer, facilitating the separation of the DNA oligos using streptavidin-functionalized magnetic beads (schematic in **Supplementary Fig. 11**).

We used this separation method to assess the strand recovery percentage after pulling out the DNA nanoaggregates.^6^ Specifically, we hybridized each oligo, in both ssDNA (ssS1 or ssS2) and ovDNA (ovS1 or ovS2) form, with its associated biotin-labeled primer, thereby generating biotin-labeled ssDNA and ovDNA (bioS1 or bioS2). We then separately incubated both biotin-labeled and unlabeled oligos (ssDNA and ovDNA) in an ionic condition containing 1 M NaCl for 20 minutes. This step resulted in the formation of individual nanoaggregates for each oligo (S1^Agg^) and (S2^Agg^) for both ssDNA and ovDNA (**Fig. 5a**). Next, we mixed the two individual nanoaggregates and incubated them for 20 minutes with only one biotin-labeled oligo. We separated each oligo using streptavidin-functionalized magnetic beads and quantified the amount of DNA for each oligo in the elution medium using qPCR.^6^ Since the streptavidin-functionalized magnetic beads were selective towards biotin-labeled oligos, we hypothesized that only the biotin-labeled DNA would be detectable in the qPCR. As expected, we observed a significant amount of DNA corresponding to biotin-labeled oligos in the elution, and a minimal quantity of DNA corresponding to the unlabeled oligos (**Fig. 5a**). This result indicated that the DNA strands can be selectively separated using consensus oligos after forming nanoaggregate structure.

**Fig. 5.**
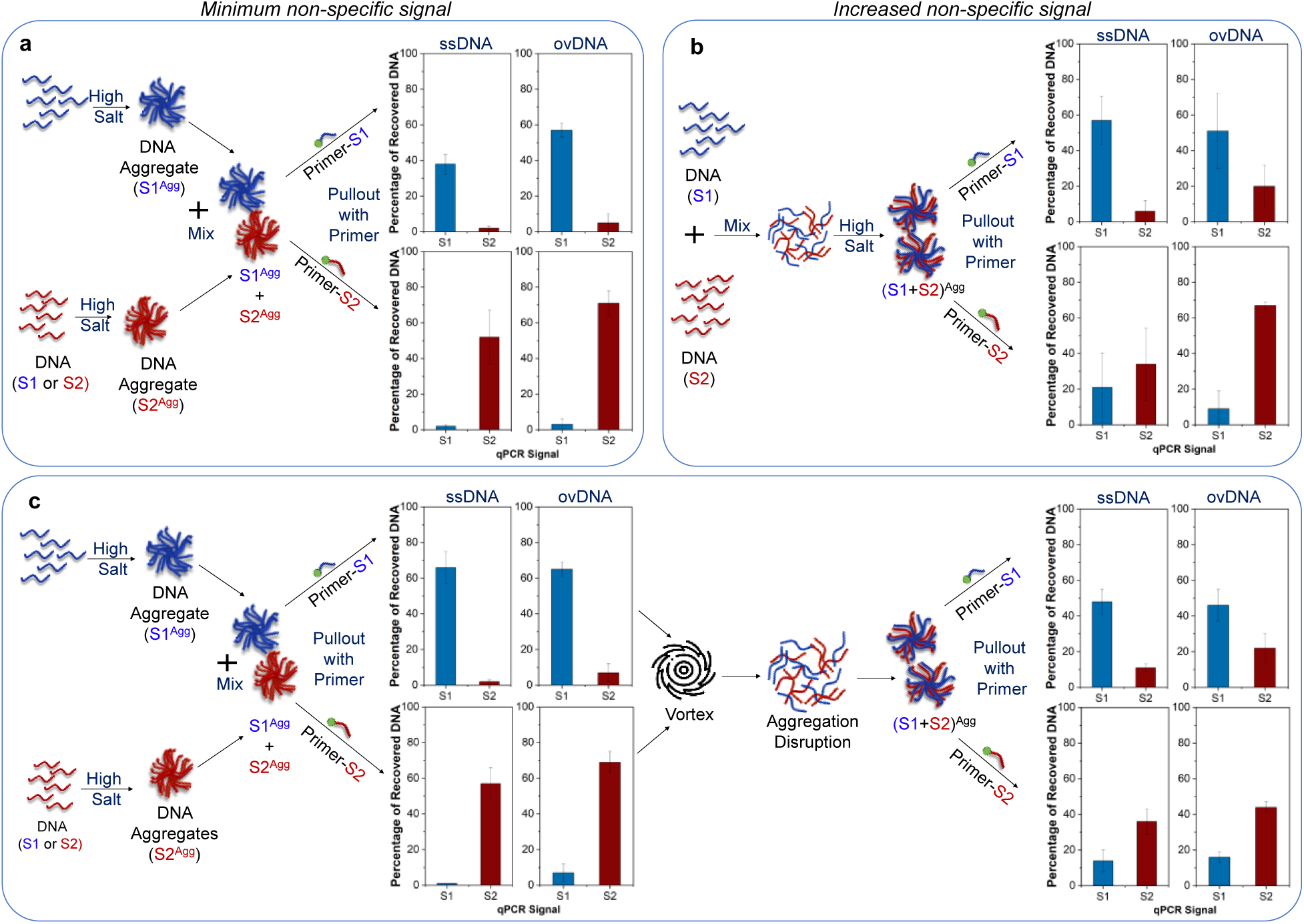
Characterization of DNA nanoaggregates using single oligos. **(a)** Schematic illustration of the formation of nanoaggregates by mixing two DNA oligos after separately incubating them for 20 minutes in 1 M NaCl. The homogeneous aggregates mixture is subjected to a pullout test using Primer-S1 and Primer-S2. DNA is quantified in the elution for each oligo. Percentage of recovery is the ratio of the amount of DNA in the elution to its original quantity. **(b)** Schematic illustration of the formation of nanoaggregates by mixing two different DNA oligos onset followed by incubating them at room temperature for 20 minutes in 1M NaCl. The heterogeneous aggregates mixture is subjected to a pullout test using Primer-S1 and Primer-S2. DNA is quantified in the elution for each oligo. **c,** Schematic illustration of the role of applied external shearing forces to previously assembled individual nanoaggregates. The aggregates mixture is subjected to a pullout test using Primer-S1 and Primer-S2. Quantification of DNA in the elution for each oligo between samples with vortexed and no-vortexed treatments.

Next, we investigated whether the selective separation can be maintained in a bundle format after assembling nanoaggregates from DNA with different sequences. We mixed S1 and S2 first and incubated the mixture in 1M NaCl solution for 20 minutes (**Fig. 5b**). We then separated the mixed DNA nanoaggregates by hybridizing biotinylated oligos that are specific to S1 or S2, followed by pulling out with streptavidin-functionalized magnetic beads, and measured the DNA quantity for each oligo using qPCR. We observed an increase in the fraction of DNA in the form of non-specific oligos pulled out in aggregates containing the specific oligos (**Fig. 5b**). Thus, the nanoaggregates can be captured and separated as intact bundles, even when they are formed of different DNA sequences.

Given that the self-assembly of DNA nanoaggregates is spontaneous, we hypothesized that the aggregation could be reversible. Thus, if the aggregates were to disassemble and subsequently reform, the resulting nanostructures might exhibit altered strand composition compared to the original aggregates. (**Fig. 3**). We start by forming individual aggregates similar to the process in **Fig. 5a**, separate each oligo using streptavidin-functionalized magnetic beads, and quantify the corresponding amount of DNA in the elution fluid using qPCR. We observed that the individual nanoaggregates preserved their composition, leading to highly specific separation. After this, we mixed the two individual nanoaggregates, incubated them at room temperature for 20 minutes, and applied external shearing forces by vortexing the mixture, followed by incubation at room temperature (**Fig. 5c**). We performed the separation of the mixture using magnetic beads and quantified the DNA amount for each oligo. Surprisingly, we observed an increase in DNA amount for non-specific oligos and a reduction in DNA amount for specific oligos (**Fig. 5c**). Taken together, these results demonstrated that while the nanoaggregates can be reformed after applying external shearing forces, their strands composition has been changed in the newly formed nanoaggregates, leading to cross-aggregation between bundles.

### Reformation of nanoaggregates can promote file scrambling in DNA-based information system

In most DNA-based storage systems^2,4–7,39–42^, primer sequences (20 nt) are included in the strand design with distinct sequences for each file to allow specific information access and separation^5^. However, the addition of these sequences significantly reduces physical space for storing digital information in DNA, thereby reducing the system capacity^5,43^. Based on the observations in nanoaggregate separation, we first sought to establish whether the file strands could be separated from the surrounding reagents when captured within nanoaggregates formed with biotin-labeled single oligos (bio-ssDNA or bio-ovDNA). We used 1174 250 nt DNA oligos, which comprised two encoded digital JPEG files (File-1: zebrafish embryo, 667 oligos; File-2: muscle cell, 507 oligos). We created dsDNA for each file strand (File-1 or File-2 DNA) and separately mixed each group of file DNA with its group of biotin-labeled pullout oligos (**Fig. 6a and b**). For example, File-1 was mixed with biotin-labeled S1 (File-1+S1), and File-2 was mixed with biotin-labeled S2 (File-2+S2). To investigate whether the ratio of the two DNAs in the nanoaggregates could influence the separation performance, we included varying ratio of pullout-oligo to file DNA, ranging from 5:1, 1:1 and 0.2:1.

**Fig. 6.**
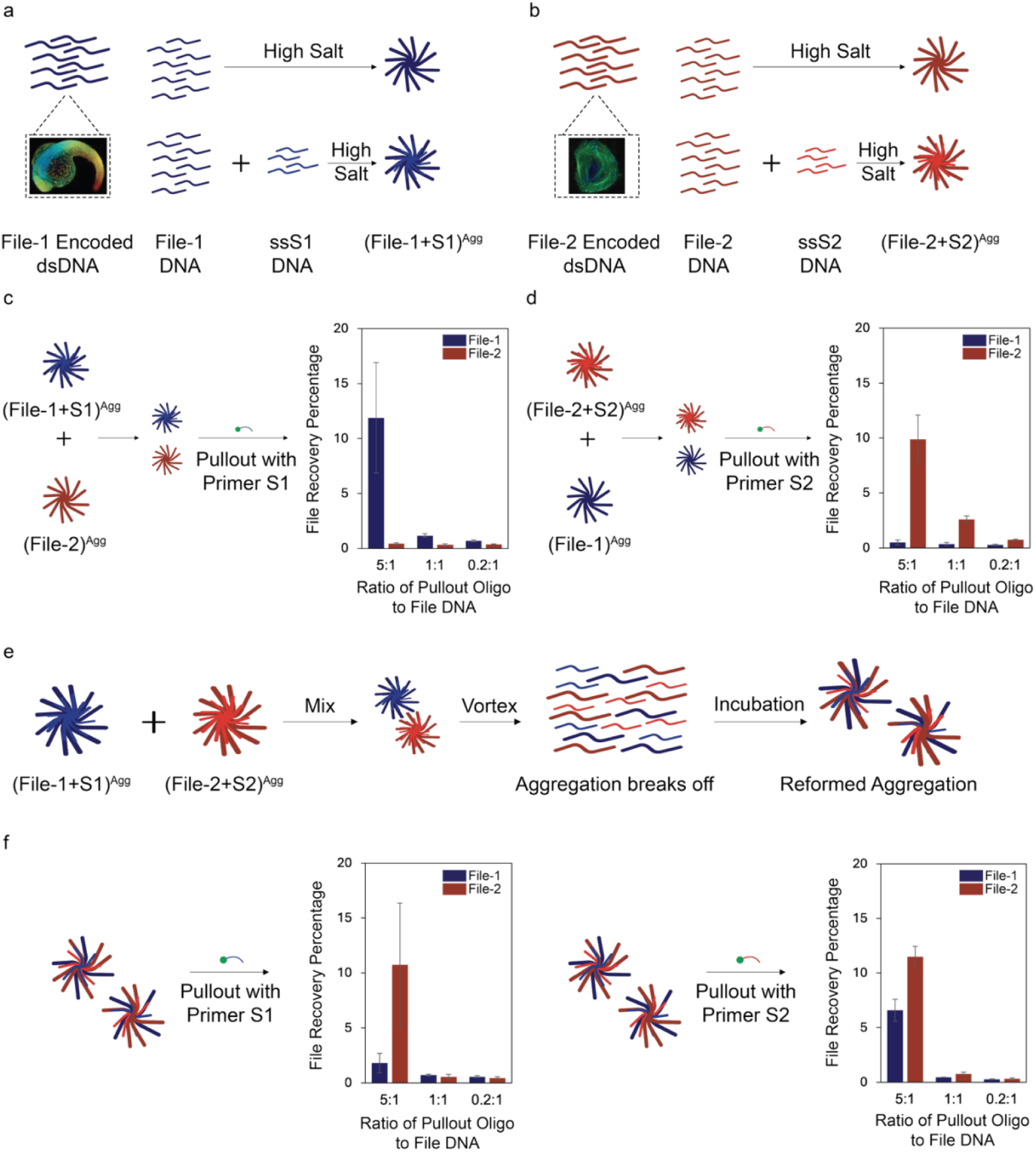
Characterization of nanoaggregates formed using file-encoded DNA after mixing with pullout oligos with and without vortexing treatments. **(a)** Schematic illustration of file-1 encoded DNA forming nanoaggregates with pullout oligos. **(b)** Schematic illustration of file-2 encoded DNA forming nanoaggregates with pullout oligos. **(c), (d)** Quantification of DNA for each file-encoded DNA in elution with varying ratios of pullout-oligo: File-encoded-DNA, ranging from 5:1, 1:1 and 0.2:1 (see Supplementary Table 2 for annotation of experimental conditions). The pullout oligos are S1 and S2, both are ssDNAs and quantifications are for File-1 and File-2 respectively. **(e)** Schematics of microshearing via vortexing of the previously assembled individual nanoaggregate containing file-encoded DNA. It results in reformed aggregates containing both File-1 and File-2. **(f)** Quantification of DNA for each file-encoded DNA in elution with varying ratios of pullout-oligo: File-encoded-DNA, ranging from 5:1, 1:1 and 0.2:1 (see Supplementary Table 3 for annotation of experimental conditions). The pullout oligos are S1 and S2, both are ssDNAs and quantifications are for File-1 and File-2 respectively. Results with pullout oligos ovS1 and ovS2 (both ovDNAs) are shown in **Supplementary Fig. 12**. The ratio of 5:1 yielded the best recovery for ovDNA as well.

All mixtures were incubated in 1M NaCl for 20 minutes to form nanoaggregates. In separate groups, we created individual nanoaggregates for the other file DNA without mixing with pullout oligo, incubating them at room temperature for 20 min (File-1)^Agg^ or (File-2)^Agg^. We assembled four sets of individual nanoaggregates: (File-1)^Agg^, (File-2)^Agg^, (File-1 + S1)^Agg^ and (File-2 + S2)^Agg^. We then mixed the two nanoaggregates together and incubated them at room temperature for another 20 min ((File-1 + S1)^Agg^ + (File-2)^Agg^) and (File-2 + S2)^Agg^ +(File-1)^Agg^). Subsequently, we performed a separation process using streptavidin-functionalized magnetic beads and quantified the DNA amount for each file strand in the elution by real-time PCR. As expected, we observed a major amount of file DNA from nanoaggregates containing pullout oligos, and a minimal quantity of file DNA from nanoaggregates that were not assembled with pullout oligos (**Fig. 6c and d**).

Next, we evaluated the strand distribution when the nanoaggregates were disrupted and reconstructed. Specifically, we applied shearing disassembly by vortexing the mixture of files- (File-1 + S1)^Agg^ and (File-2 + S2)^Agg^, for 5 minutes, followed by incubation for 20 mins (**Fig. 6e**). We then performed the separation process and quantified the amount of DNA for each file strand in the elution medium. Excitingly, we observed an increase in the DNA amount for the file DNA that was not assembled with pullout oligos, and a decrease in the DNA amount for the file DNA that was assembled with pullout oligos (**Fig. 6f**). This suggested that the previously aggregated file strands, which neatly comprised individual file information, were scrambled after disrupting and reforming the nanoaggregates. File recovery results from experiments where the aggregates were formed using ovDNA are presented in **Supplementary Fig. 12**.

An assembly ratio of 5:1 between pullout oligo and file DNA resulted in the highest DNA recovery rate in the elution fluid (**Supplementary Fig. 13** and **Supplementary Table 4**) and was therefore selected for next-generation sequencing (NGS) analysis. Additional studies using alternative ratios of pullout oligo to file DNA (1:0.2, 1:1, and 1:5; **Supplementary Fig. 14**) produced lower recovery rates compared to the 5:1 ratio. Elution samples from this ratio were prepared for NGS analysis and confirmed the strand scrambling in reassembled nanoaggregates (**Fig. 7a and b**). An important observation is that strands scrambling was encountered in a relatively low amount. This could be due to the degree of persistence of dsDNA when forming the nanostructure with other single or double-stranded oligos.

**Fig. 7.**
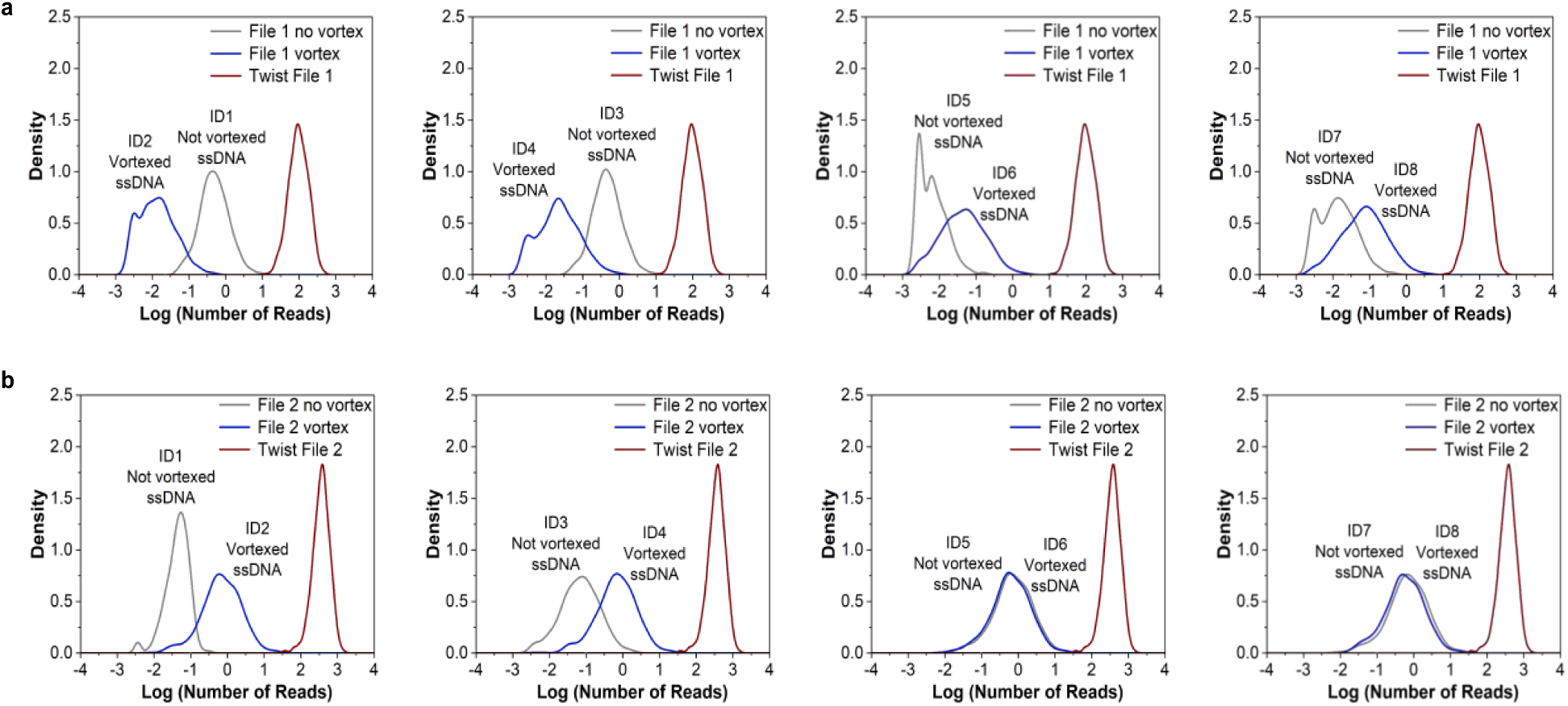
NGS confirms strand scrambling for reconstructed nanoaggregates. **(a)** Density plot displaying the strand distribution of samples containing “Agg(bioS1+file1) + Agg(file2)” with vortexing treatment, compared to the same samples without vortexing treatment. The plot includes file-encoded strands passed through File-1 decoding algorithm (see Supplementary Table 5 for annotation of experimental condition). **(b)** Density plot displaying the strand distribution of samples containing “Agg(bioS2+file2) + Agg(file1)” with vortexing treatment, compared to the same samples without vortexing treatment. The plot includes file-encoded strands passed through File-2 decoding algorithm (see Supplementary Table 6 for annotation of experimental conditions).

In summary, nanoaggregates formed by single DNA oligos could serve as file addresses to assemble in nanoaggregates with a pool of DNA encoding digital information, enabling the separation of targeted information under optimized ionic conditions. However, the information encoded in two types of nanoaggregates can be exchanged between them by disrupting and reforming them, allowing for “mechanical” information scrambling in a DNA-based storage system.

### Understanding Nanoaggregation Dynamics for DNA Data Storage: From Assembly to Disruption

The DNA assemblies are formed as a result of DNA-DNA interactions in the presence of electrolytes. DNA molecules have negatively charged phosphate strands along their backbone. Positive ions such as Na^+^ can bind to the major and minor grooves of DNA. The counterions form "ion clouds" around the negatively charged phosphate groups, suppressing the electrostatic repulsion between them^19^. At low salt concentrations, the charge effects in polyelectrolyte chains such as DNA are described by the Debye-Hückel theory. The Debye length (*1/κ*) is related to the ionic strength (*I*) as 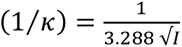. The ionic strength depends on the concentration of electrolyte and the ion valency 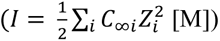. At low ionic strength, the Debye length is relatively high and leads to long-range electrostatic interactions. However, it plunges at higher ionic strengths, e.g. 1 M CaCl_2_ reduces it to 0.17 nm. The strong electrostatic shielding reduces the inter-strand repulsion and induces aggregation. Additionally, the suppression of the electrostatic repulsion reduces the persistent length, which makes the chains more flexible^44^. More flexible chains can pack more densely, which could be the reason for the self-assembly of the DNA strands as observed in **Fig. 2a**. dsDNA has a propensity to form aggregates sooner owing to the greater charge density of dsDNA^45^. Our data in **Fig. 2a** support these expectations.

**Fig. 8a** illustrates two possible mechanisms behind the formation of these aggregates in the presence of Na^+^. Charged chains such as DNA strands attract each other in the presence of counterions (inter-DNA attractions)^46–49^. DNA is highly polyvalent which facilitates ion bridging. The negative charges of the phosphates along the length of the strands may interact with a “zipper” mechanism which pulls multiple strands together^50–52^. The combined effect of DNA electrostatic and depletion-like attractions has been calculated by several molecular dynamic simulations^22,53–55^.

**Fig. 8.**
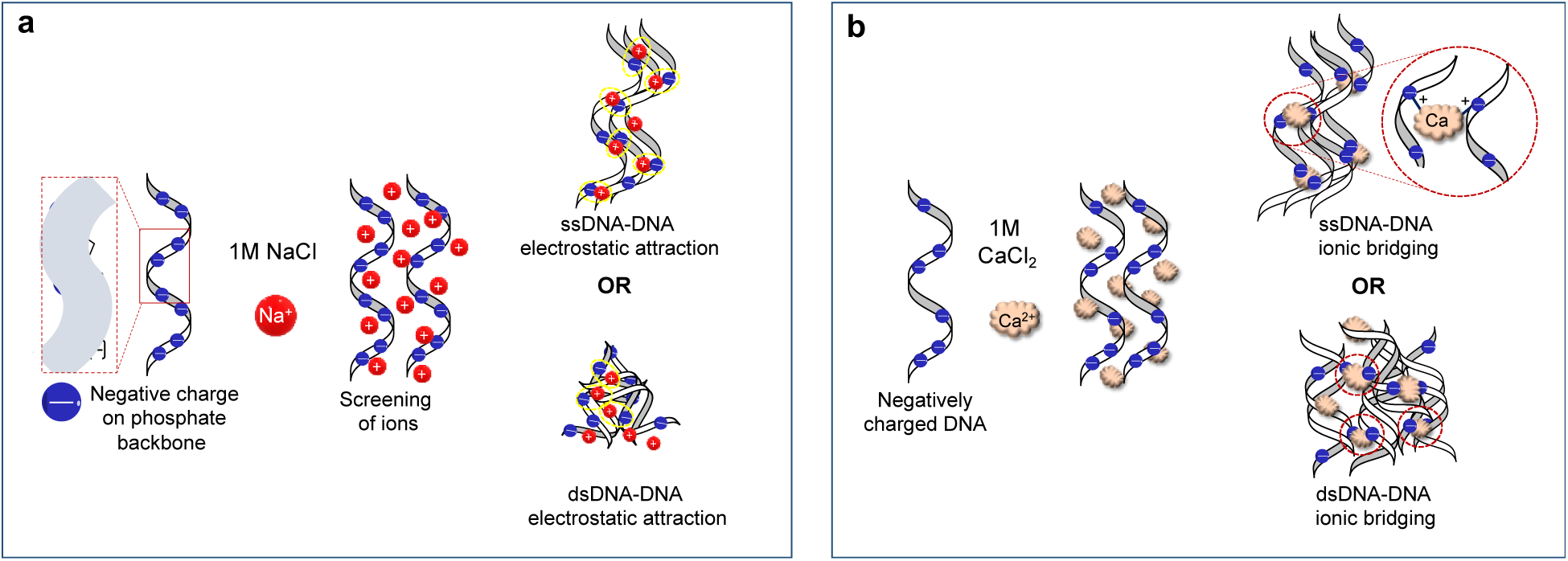
Ion-mediated associations between the DNA strands leading to the formation of nanoaggregates. **(a)** Schematic illustrating DNA-DNA attractive interactions for ssDNA and dsDNA in presence of Na^+^ ions. **(b)** Schematic illustrating the formation of ionic bridges for ssDNA and dsDNA in presence of Ca^2+^ ions.

We observed that dsDNA aggregates are more compact as compared to ssDNA aggregates (**Fig. 2b and c**). In high ionic medium, ssDNA also experiences intra-sequence self-stacking interactions amongst nearby bases^56^. Therefore, ssDNA can adopt a variety of conformations, potentially leading to the formation of larger aggregates. In contrast, the high negative charge and higher rigidity of dsDNA^57^, lead to secondary or tertiary structures that are more condensed at high salt concentrations. This is supported by our results that show a tighter size distribution indicating the formation of more condensed aggregates for dsDNA as compared to ssDNA (**Supplementary Fig. 3**). Additionally, in the case of dsDNA there is a possibility that the aggregation is aided by the presence of a residual protein (Q5 DNA polymerase) which is used during the synthesis of dsDNA. Overall, the nanoaggregates, including the lyophilized ones (**Fig. 2d**), formed at higher ionic concentrations are thermodynamically favorable^16^ and are revealed here to remain stable over extended periods.

The molecular differences between DNA and RNA leads to subtle differences in the size and structure of the nanoaggregates. In **Fig. 2e**, the variation in GC content did not affect the nature of the aggregates. As expected, there is no significant difference in the nature of secondary structures and the electrostatic attractions are governed mainly by the charge neutralization of phosphates; therefore, the aggregates do not vary much in size.^58–60^ However, the size of aggregates of RNA where Na^+^ ions neutralize the backbone charge repulsion is smaller compared to ssDNA and is not as well-defined (**Fig. 2f**). This can be attributed to the fact that RNA is not as structurally stable and has the tendency to form multiple secondary and tertiary structures (such as loops, hairpins and bulges). Salt ions in the solution facilitate the reorganization and formation of such structures.^61^

Increasing the ionic strength and counterion valency enhances short-range attractive interactions, making them more pronounced. **Fig 8b** explains the mechanism of interactions between the strands in the presence of Ca^2+^. Divalent cations such as Ca^2+^ or Mg^2+^ also bring higher electrostatic shielding as compared to Na^+^ due to the “bridge effect” between two negatively charged adjacent DNAs.^19^ Moreover, Ca^2+^ is proven to overwind DNA significantly more than any other mono- or divalent ion,^17^ justifying the higher size of these densely condensed aggregates.

We observed reversible aggregates for 2 M NaCl but irreversible aggregation with 1M CaCl_2_ in our dynamic aggregation studies (**Fig. 3**). For relatively low ionic strengths (I ≤ 2M), the shear forces generated by vortexing can overcome the attractive forces which are loosely bounded and are primarily weak and disperse the aggregates. When the sample is measured at a later time point after vortexing, the plot indicates reaggregation. Contrastingly, no major change occurred on vortexing for the DNA aggregates formed at higher ionic strength (I = 3M). This leads to highly condensed and stable DNA aggregates,^58^ that are more resistant to shear-induced breakup. Moreover, divalent cations (Ca^2+^) likely create strong bonds between the DNA strands owing to ionic bridging^62^.

The nanobundles and nanoclusters observed in TEM micrograph validate the concept of ion-mediated DNA-DNA aggregation in well-defined clusters. Positively charged ions in solution bind to DNA and increase its flexibility as a larger fraction of the negatively charged phosphate groups would be neutralized. Zipper mechanism pulls different strands together and these interbound strands of DNA form distinct, defined “nanobundles” as seen in **Fig. 4a and b**. The nanoclusters could be a result of multiple dsDNA strands coming together and stabilizing in the presence of monovalent cations. dsDNA has larger charge density and therefore has the propensity to form more condensed structures. The presence of minor amounts of DNA polymerase in this solution may also result in more condensed packing of dsDNA strands. Similar structures with distinct toroidal shape have been reported earlier^63,64^.

Our results demonstrated that the nanoaggregates formed from distinct DNA oligos maintained their nanostructure integrity. One potential application of this phenomenon in a DNA-based storage system is the grouping of archival data files. Aggregation of file-encoded DNA oligos with non-encoded oligos resulted in the formation of individual nanostructures, where the non-encoded oligos served as file addresses in kind for the nanoaggregates, which thanks to them may be specifically separated to access the included information. This finding has significant implications for the development of a DNA-based storage systems, where two digital files were encoded in DNA sequences. By mixing non-encoded DNA oligos with file-encoded oligos, the non-encoded oligos can serve as the unified file address for each grouped information, and specific groups of information can be accessed by separation of its non-encoded oligos. This grouping and information accessing process is similar to partitioning file-encoded DNA into multiple packets.^65^ As each packet contains the group information, separation and data management are significantly simplified, compared to accessing each individual file by its distinct file address in the current systems.

Interestingly, file-encoded oligos from previously assembled individual nanoaggregates could be exchanged and their information was scrambled when external shearing forces were applied, leading to the disruption and reconstruction of the nanoaggregates. After applying of external shear forces, the strand scrambling phenomenon can be used for data obfuscation in a data storage system, in which targeted information designed for specific access can be randomly mixed within the noisy background DNA information when certain conditions are triggered^66,67^. This offers an additional layer of data security for data stored as DNA sequence strands, which become mixed up and hidden within aggregates of non-sequential DNA.

## Conclusions

We characterize and evaluate the potential for use in information storage of DNA nanoaggregates formed in high salinity water media. We found that when subjected to high ionic concentrations, molecular DNA forms stable, well-defined and reproducible nanoaggregates. High concentration of ions in the solution leads to self-assembly between the DNA strands and formation of the nanoaggregates. These nanoaggregates are found to be stable for days without any precipitation. Variations in the GC content in DNA sequences does not influence the assembly of aggregates. It was noticed that there is a difference in the nanostructures for ss and dsDNA. ssDNA forms “nanobundles” or “rod-like” structures whereas dsDNA form “nanoclusters” in high salt concentration solutions. Similar behavior is observed with RNA, but the aggregates are less stable. Salts of divalent ions (Ca^2+^) form larger sized aggregates due to the bridging effect and greater electrostatic shielding.

Given the mild conditions of DNA nanoaggregate self-assembly, we hypothesized that their formation might be reversible. When the system was subjected to fluid shear forces, we observed that the aggregates dissociated into smaller entities, and reassembled after the system was returned to rest. However, in the presence of divalent ions these aggregates show a much smaller tendency to disintegrate under shear. Lower ionic strength allows reversible aggregation, whereas higher ionic strength results in irreversible aggregation.

This system offers several avenues for further advancement. These nanoaggregates can also be functionalized to attach fluorescent molecules to enhance visualization. Further optimization of the ratio of non-encoded oligo and file-encoded DNA and refinement of separation conditions can increase the yield of recovered file-encoded DNA in the elution. It is also important to investigate how different lengths of non-encoded oligos affect file-encoded DNA and their impact on file recovery. Overall, our findings highlight that DNA aggregates can be used to enhance the efficacy of DNA-based storage systems. Understanding the self-assembly phenomena in DNA nanostructures opens up new possibilities for achieving efficient and secure information storage at high densities. Further research and development in this field hold promise for innovative methods for data storage and encryption.

### 4.2 Experimental Methods

#### 4.2.1. Formation of the Nanoaggregates

ssDNA(120nt) oligos were purchased from Eton Bioscience Inc. which was diluted in filtered (0.2μm syringe filter Millipore Sigma) DI water (Millipore-Q, resistivity 18.2 MΩ cm) to a concentration of 100 μM. Small aliquots were made for subsequent uses and the sample was stores in -20°C. Each aliquot was thawed and diluted to the desired concentration of DNA. ssDNA (200nt) oligos was purchased from Integrated DNA Technologies (IDT) Inc. and diluted to a concentration of 10μM and made into aliquots. 120nt ssDNA with 0, 25, 50 and 75% of G-C in their sequences was ordered from IDT Inc and was diluted to 10μM as well. Their specific secondary structures were predicted by VectorBuilder Inc. Sequences are listed in supporting information in Supplementary Table 1. The sequences for the study were generated using a random sequence generator tool available online. RNA and dsDNA were created in the lab using the protocol listed below.

Solutions of 0.001M, 0.01M, 0.1M, 0.5M, 1M, 2M Sodium Chloride (extra pure, Thermo Scientific), 1M Calcium Chloride (VWR Life Science) and 1M Aluminum Chloride (Sigma) were prepared in filtered DI water. DNA aliquots were thawed and centrifuged for 5-10 mins. DNA was added to solutions of varying salt concentrations maintaining a final concentration of 0.1μM of DNA in solution. The DNA-salt suspensions were prepared in room temperature. Soon after, these suspensions were subjected to various characterizations.

#### 4.2.2. Nanoaggregates Characterizations

DNA and Nanoaggregate sizes and zeta potential were measured through dynamic light scattering using a Zetasizer Nano ZSP (Malvern Instruments Ltd.) at room temperature (25°C) and increased temperatures. The Zetasizer was fitted with a 633 nm He-Ne laser and performed measurements with a 173° backscattering mode. Each measurement included 9 or 12 subruns and data was generated in triplicates. pH of samples was measured using a Mettler-Toledo pH meter. To lyophilize the samples, two twist-top freezer tubes containing ssDNA and dsDNA in high salt solutions were frozen at -80°C overnight. The tube tops were slightly loosened, and the samples were placed in a benchtop freeze-dryer (Labconco FreeZone, 2.5L) for 21 hours at -50°C and 0.2 mbar. After lyophilization, the tube caps were tightened, and the samples were stored at -20°C. They were rehydrated with DI water before measurements.

A Biological Transmission Electron Microscope (120kV, HT7800, Hitachi, Tokyo, Japan) was used to visualize the nanoaggregates. The samples were deposited on a glow-discharged TEM grids and stained with uranyl acetate. The accelerating voltage was 80kV and the images were captures at 60× magnification. After 20 mins, excess sample was blotted off using filter paper and the grids were stained with 2% uranyl acetate solution. The grids were placed in a desiccator and visualized the next day. ImageJ (NIH, USA) software was used to determine the average size of nanoaggregates. Multiple size measurements were obtained from different TEM images for the same set of samples.

#### 4.2.3. Disruption of Aggregation and Reaggregation

The DNA-salt suspensions containing nanoaggregates were subjected to mechanical shearing by vortexing for 5 mins. The solution was kept undisturbed for 20-40 mins before taking further measurements.

#### 4.2.4. Preparation of RNA by DNA Replication

A DNA oligo with a T7 promoter sequence was purchased from Azenta, Inc and used as a template to create dsT7-DNA using PCR, followed by purification with AMPure XP beads (Beckman Coulter, A63881) and elution in 40 μL of water. 300 ng of dsT7-DNA was mixed with 30 µL of in vitro transcription buffer (NEB, E2050) containing 2 µL of T7 RNA Polymerase Mix and ATP, TTP, CTP, GTP, each at 6.6 mM. The mixture was incubated at 37 °C for 16 h and purified by a Monarch RNA 10 Cleanup Kit (NEB, T2040L) following the manufacturer’s instructions. The newly generated RNA transcripts were measured using a NanoDrop Spectrophotometer and Fragment Analyzer HS RNA Kit (Agilent Technologies Inc., DNF-472-0500).

#### 4.2.5. Preparation of ovDNA

ovDNA strands were created by filling in ssDNA templates (Integrated DNA Technologies Inc.) with primer TCTGCTCTGCACTCGTAATAC (Azenta Inc.) at a ratio of 1:40 using 0.5 µL of Q5 High-Fidelity DNA Polymerase (NEB, M0491S) in a 50 µL reaction containing 1x Q5 polymerase reaction buffer (NEB, B9072S) and 2.5 mM each of dATP (NEB, N0440S), dCTP (NEB, N0441S), dGTP (NEB, N0442S), dTTP (NEB, N0443S). The reaction conditions were 98 °C for 30 s and then 4 cycles of: 98 °C for 10 s, 53 °C (1 °C s−1 temperature drop) for 20 s, 72 °C for 10 s, with a final 72 °C extension step for 2 min. ss-dsDNA strands were purified using AMPure XP beads (Beckman Coulter, A63881) and eluted in 40 μL of water.

#### 4.2.6. Preparation of ssDNA Nanoaggregates for Characterization Using Single Oligos

ssDNA (Azenta Inc.) strands were diluted to 1011 strands and mixed with 5’ biotinylated oligos (Integrated DNA Technologies Inc.) at a ratio of 1:40 in a 30 µL reaction containing 2 mM MgCl_2_ (Invitrogen, Y02016) and 50 mM KCl (NEB, M0491S). Oligo annealing conditions were 45 °C for 2 min, followed by a temperature drop at 1 °C/min to 14 °C. The mixture was further supplemented with additional 5M NaCl to a final concentration of 1 M and a final reaction volume of 50 µL, followed by incubating at room temperature for 20 min.

#### 4.2.7. Preparation of ovDNA Nanoaggregates for Characterization Using Single Oligos

ovDNA strands were obtained from previous instructions and were diluted to 1011 strands and mixed with 5’ biotinylated oligos (Integrated DNA Technologies Inc.) at a ratio of 1:40 in a 30 µL reaction containing 2 mM MgCl_2_ (Invitrogen, Y02016) and 50 mM KCl (NEB, M0491S). Oligo annealing conditions were 45 °C for 2 min, followed by a temperature drop at 1 °C/min to 14 °C. The mixture was further supplemented with additional 5M NaCl to a final concentration of 1 M and a final reaction volume of 50 µL, followed by incubating at room temperature for 20 min.

#### 4.2.8. Creation of Nanoaggregates by Mixture for Characterization Using Single Oligos

Pre-annealed ssDNA or ovDNA strands were mixed with another sequence of ssDNA or ovDNA, respectively, in a 50 uL reaction containing 2 mM MgCl_2_ (Invitrogen, Y02016) and 30 mM KCl (NEB, M0491S). The mixture was supplemented with additional 5M NaCl to a final concentration of 1 M and a final reaction volume of 50 µL, followed by incubating at room temperature for 20 min.

#### 4.2.9. Reformation of Nanoaggregates after Disruption for Molecular Studies

Pre-assembled nanoaggregates were placed on a mini vortexer (VWR) at 2000 RPM for 5 mins. The mixture should be secured on the vortexer and maintain the upward position during the vortexing. This was followed by incubating the mixture at room temperature for 20 min.

#### 4.2.10. Separation of ss/ovDNA Nanoaggregates

Streptavidin magnetic beads (NEB, S1420S) were prewashed using high salt buffer containing 20 mM Tris-HCl, 2 M NaCl and 2 mM EDTA pH 8 and incubated with nanoaggregates pre-annealed with biotinylated oligos at room temperature for 30 min. The supernatant was collected for downstream real-time PCR analysis. The beads were washed with 100 µL of high salt buffer and subsequently eluted with 95% formamide (Sigma, F9037) in water. The quality and quantity of the DNA eluted from the beads were measured by real-time PCR (Bio-Rad).

#### 4.2.11. File Encoding and Decoding Designs for Zebrafish Embryo and Muscle Cell

The file DNA was obtained from previous research. Briefly, two digital files were partitioned into blocks of data that fit in DNA strands with 250 nt length. Each strand consisted of multiple sections. A primer binding sites were positioned at each end for DNA amplification using PCR. Following the 5’ end primer site, a synthetic T7 promoter sequence and second primer site were incorporated. The second primer site can allow the amplification of complementary DNA (cDNA) after reverse transcription (RT). In between the second 5’ primer site and 3’ primer site, digital file information was encoded and partitioned into three sections including the index of the strand within the file, the data payload, and a checksum to detect errors within the strand. We designed 8 bp-long codewords to represent one byte of data. The codewords have no repetition of bases both individually and when appended, and they are GC balanced. Each byte of file is converted one byte at a time into a corresponding codeword and appended together to form the payload of a strand. We also adopted a redundant XOR-style encoding proposed by Bornholt et al. to enhance the reliability of our system. The decoder algorithm for our encoding is similar to that used in previous work with the modification that we can disregard any read with an invalid checksum.

#### 4.2.12. Primer Design

Primer used in this worked were designed based on the following criteria: 1) GC content is between 40 and 60%; 2) their melting temperature is between 50 and 60 °C; 3) the last base is G, but the GC content in the last 5 bases could not exceed 60% and 4) no heterodimer bindings. Meanwhile, primers were designed to reduce the likelihood of nonspecific binding with other primer binding sites. Hamming distance of >10 between all primers was required to minimize the likelihood of such binding. We also required a Gibbs free energy greater than −10 kcal/mol at 50 °C on all likely complexes to select the primer.

#### 4.2.13. Preparation of dsDNA (file DNA)

ssDNA library that contained digital information were designed and purchased from Twist Bioscience Inc. Unlabeled primer oligos were purchased from Azenta Inc. 0.1 ng of ssDNA was mixed with unlabeled primer oligos at a final concentration of 1 µM using 0.5 µL of Q5 High-Fidelity DNA Polymerase (NEB, M0491S) in a 50 µL reaction containing 1x Q5 polymerase reaction buffer (NEB, B9072S) and 0.2 mM each of dATP (NEB, N0440S), dCTP (NEB, N0441S), dGTP (NEB, N0442S), dTTP (NEB, N0443S). The reaction conditions were 98 °C for 30 s and then 35 cycles of: 98 °C for 10 s, 53 °C for 20 s, 72 °C for 10 s, with a final 72 °C extension step for 2 min. The generated unlabeled dsDNA strands were examined using gel electrophoresis and purified using Monarch Gel Extraction Kit (NEB, T1020S) following manufacturer’s instructions.

#### 4.2.14. Gel electrophoresis of DNA samples

Agarose-based DNA gels were made by mixing and microwaving 100 mL of 1x TAE buffer (Fisher Scientific, BP13324) with 1.5 mg of molecular biology grade agarose (Genesee Scientific, 20102). 0.1x SYBR Safe DNA Gel Stain was added to visualize DNA (Invitrogen, S33102). DNA samples and ladder (NEB, N3231S) were loaded with 1x DNA loading dye containing 10 mM EDTA, 3.3 mM Tris-HCl (pH 8.0), 0.08% SDS and 0.02% Dye 1 and 0.0008% Dye 2 (NEB, B7024S). Electrophoresis was performed with 1x TAE buffer in a Thermo Scientific Mini Gel Electrophoresis System (Fisher Scientific, 09–528–110B) at a voltage gradient of 16 V/cm for 45 min. Purification of DNA in the gel was achieved by using Monarch Gel Extraction Kit (NEB, T1020S) following manufacturer’s instructions.

#### 4.2.15. Gel Imaging

Fluorescence imaging of both DNA and RNA gel samples was performed with a Li-Cor Odyssey® Fc Imaging System and the fluorescence intensity was quantified using FIJI software.

#### 4.2.16. Preparation of File DNA Nanoaggregates with Pullout DNA

Pullout DNA strands (ssDNA or ovDNA) were pre-annealed with biotinylated oligos using previous instructions. The mixture was then added with file DNA (dsDNA) at a ratio of 5:1, 1:1 and 0.2:1 or 1:0.2, 1:1 and 1:5 in a 30 µL reaction containing 2 mM MgCl_2_ (Invitrogen, Y02016) and 50 mM KCl (NEB, M0491S). Oligo annealing conditions were 45 °C for 2 min, followed by a temperature drop at 1 °C/min to 14 °C. The mixture was further supplemented with additional 5M NaCl to a final concentration of 1 M and a final reaction volume of 50 µL, followed by incubating at room temperature for 20 min.

#### 4.2.17. Real-time PCR (qPCR)

qPCR was performed in a 6 μL, 384-well plate format using SsoAdvanced Universal SYBR Green Supermix (BioRad, 1725270). The amplification conditions were 95 °C for 2 min and then 50 cycles of: 95 °C for 15 s, 53 °C for 20 s, and 60 °C for 20 s. Quantities were interpolated from the linear ranges of standard curves performed on the same qPCR plate.

#### 4.2.18. Next-generation Sequencing

Amplicons were purified with AMPure XP beads (Beckman Coulter, A63881) according to the TruSeq Nano protocol (Illumina, 20015965). The quality and band sizes of libraries were assessed using the HS NGS Fragment Analysis Kit (Advanced Analytical, DNF-474) on the 12 capillary Fragment Analyzer (Agilent Technologies Inc.). The prepared samples were submitted to Azenta Inc. for Illumina-based next-generation sequencing (Amplicon-EZ). Ligation of Illumina sequencing adapters to the prepared samples was performed by Azenta Inc. Data analysis was performed by using FLASH v1.2.11 from Conda v23.5 for QC and using Pandas v2.0.2 from Python v3.8 to sort the number of reads for each strand.

## Acknowledgments

The authors thank Prof. Robert M. Kelly for the use of his vacuum concentrator and appreciate the assistance of Lauren Kielty and Tyler Wroblewski with some experiments, as well as Dr. Anuj Kumar for Python-based data analytics. They also thank Karishma Matange for her help with the lyophilizer. This work was performed in part at the Analytical Instrumentation Facility (AIF) at North Carolina State University, which is supported by the State of North Carolina and the National Science Foundation (award number ECCS-2025064). The AIF is a member of the North Carolina Research Triangle Nanotechnology Network (RTNN), a site in the National Nanotechnology Coordinated Infrastructure (NNCI). We are thankful to Dr. Aaron Bell and Dr. Chris Winkler at AIF, NCSU.

## Funding

This work was supported by the US National Science Foundation (ECCS-2027655 and partially DMR-2303581 and DMR-2243104). KNL was supported by a Department of Education Graduate Assistance in Areas of Need fellowship.

## Author contributions

Conceptualization: SM, KNL, AJK, ODV

Methodology: SM, KNL, AJK, ODV

Investigation: SM, KNL, KV

Visualization: SM

Supervision: JMT, AJK, ODV

Writing—original draft: SM, KNL

Writing—review & editing: SM, KNL, AJK, ODV

## Competing interests

Authors declare that they have no competing interests

## Data and materials availability

All data are available in the main text or the supplementary materials.

## Notes

### Competing Interest Statement

The authors have declared no competing interest.

## References

1. Meiser, L. C. et al. Synthetic DNA applications in information technology. Nat Commun 13, 352 (2022).

2. Goldman, N. et al. Towards practical, high-capacity, low-maintenance information storage in synthesized DNA. Nature 494, 77–80 (2013).

3. Church, G. M., Gao, Y. & Kosuri, S. Next-generation digital information storage in DNA. Science 337, 1628 (2012).

4. Bornholt, J. et al. A DNA-Based Archival Storage System. in Proceedings of the Twenty-First International Conference on Architectural Support for Programming Languages and Operating Systems 637–649 (Association for Computing Machinery, New York, NY, USA, 2016). doi:10.1145/2872362.2872397.

5. Organick, L. et al. Random access in large-scale DNA data storage. Nat Biotechnol 36, 242–248 (2018).

6. Lin, K. N., Volkel, K., Tuck, J. M. & Keung, A. J. Dynamic and scalable DNA-based information storage. Nat Commun 11, 2981 (2020).

7. Lin, K. N. et al. A primordial DNA store and compute engine. Nat. Nanotechnol. 1–11 (2024) doi:10.1038/s41565-024-01771-6.

8. Tomek, K. J. et al. Driving the Scalability of DNA-Based Information Storage Systems. ACS Synth Biol 8, 1241–1248 (2019).

9. Mills, A., Aissaoui, N., Finkel, J., Elezgaray, J. & Bellot, G. Mechanical DNA Origami to Investigate Biological Systems. Advanced Biology 7, 2200224 (2023).

10. Li, R., Madhvacharyula, A. S., Du, Y., Adepu, H. K. & Choi, J. H. Mechanics of dynamic and deformable DNA nanostructures. Chem Sci 14, 8018–8046 (2023).

11. Harmouchi, M., Albiser, G. & Premilat, S. Effect of a mechanical tension on the hydration of DNA in fibres. Biochemical and Biophysical Research Communications 188, 78–85 (1992).

12. Burak, Y., Ariel, G. & Andelman, D. Onset of DNA Aggregation in Presence of Monovalent and Multivalent Counterions. Biophys J 85, 2100–2110 (2003).

13. Duguid, J. G. & Bloomfield, V. A. Electrostatic effects on the stability of condensed DNA in the presence of divalent cations. Biophys J 70, 2838–2846 (1996).

14. Soumpasis, D. M. Salt dependence of DNA structural stabilities in solution. Theoretical predictions versus experiments. J Biomol Struct Dyn 6, 563–574 (1988).

15. Soumpasis, D. M., Wiechen, J. & Jovin, T. M. Relative stabilities and transitions of DNA conformations in 1:1 electrolytes: a theoretical study. J Biomol Struct Dyn 4, 535–552 (1987).

16. Tan, Z.-J. & Chen, S.-J. Nucleic Acid Helix Stability: Effects of Salt Concentration, Cation Valence and Size, and Chain Length. Biophys J 90, 1175–1190 (2006).

17. Cruz-León, S. et al. Twisting DNA by salt. Nucleic Acids Research 50, 5726–5738 (2022).

18. Zavadlav, J., Podgornik, R. & Praprotnik, M. Adaptive Resolution Simulation of a DNA Molecule in Salt Solution. J. Chem. Theory Comput. 11, 5035–5044 (2015).

19. Yu, Q., Chen, J., Shi, D. & Chen, M. Ordered self-assembly of DNA-modified nanoparticles in salt solutions. Colloids and Surfaces A: Physicochemical and Engineering Aspects 671, 131669 (2023).

20. Zhang, Z. et al. Salt-Induced Assembly Transformation of DNA–AuNP Conjugates Based on RCA Origami: From Linear Arrays to Nanorings. Langmuir 34, 8904–8909 (2018).

21. Lee, L., Cavalieri, F., Johnston, A. P. R. & Caruso, F. Influence of Salt Concentration on the Assembly of DNA Multilayer Films. Langmuir 26, 3415–3422 (2010).

22. Seo, S. E., Girard, M., de la Cruz, M. O. & Mirkin, C. A. The Importance of Salt-Enhanced Electrostatic Repulsion in Colloidal Crystal Engineering with DNA. ACS Cent Sci 5, 186–191 (2019).

23. Oh, J.-H. & Lee, J.-S. Salt concentration-induced dehybridisation of DNA–gold nanoparticle conjugate assemblies for diagnostic applications. Chem. Commun. 46, 6382–6384 (2010).

24. DeRouchey, J. E. & Rau, D. C. Salt Effects On Condensed Protamine-Dna Assemblies: Anion Binding And Weakening Of Attraction. J Phys Chem B 115, 11888–11894 (2011).

25. Misra, V. K., Hecht, J. L., Sharp, K. A., Friedman, R. A. & Honig, B. Salt Effects on Protein-DNA Interactions: The λcI Repressor and EcoRI Endonuclease. Journal of Molecular Biology 238, 264–280 (1994).

26. Ross, M. B., Ku, J. C., Vaccarezza, V. M., Schatz, G. C. & Mirkin, C. A. Nanoscale form dictates mesoscale function in plasmonic DNA–nanoparticle superlattices. Nature Nanotech 10, 453–458 (2015).

27. Hanke, M. et al. Salting-Out of DNA Origami Nanostructures by Ammonium Sulfate. Int J Mol Sci 23, 2817 (2022).

28. Hübner, K., Raab, M., Bohlen, J., Bauer, J. & Tinnefeld, P. Salt-induced conformational switching of a flat rectangular DNA origami structure. Nanoscale 14, 7898–7905 (2022).

29. Zhang, F., Nangreave, J., Liu, Y. & Yan, H. Structural DNA Nanotechnology: State of the Art and Future Perspective. J. Am. Chem. Soc. 136, 11198–11211 (2014).

30. Endo, M. & Sugiyama, H. DNA Origami Nanomachines. Molecules 23, 1766 (2018).

31. Lacroix, A. & Sleiman, H. F. DNA Nanostructures: Current Challenges and Opportunities for Cellular Delivery. ACS Nano 15, 3631–3645 (2021).

32. Ma, W. et al. The biological applications of DNA nanomaterials: current challenges and future directions. Sig Transduct Target Ther 6, 1–28 (2021).

33. Ganesh, A. N., Donders, E. N., Shoichet, B. K. & Shoichet, M. S. Colloidal aggregation: From screening nuisance to formulation nuance. Nano Today 19, 188–200 (2018).

34. Scipioni, A., Anselmi, C., Zuccheri, G., Samori, B. & Santis, P. D. Sequence-Dependent DNA Curvature and Flexibility from Scanning Force Microscopy Images. Biophysical Journal 83, 2408–2418 (2002).

35. Merivaara, A. et al. Preservation of biomaterials and cells by freeze-drying: Change of paradigm. Journal of Controlled Release 336, 480–498 (2021).

36. Anchordoquy, T. J., Carpenter, J. F. & Kroll, D. J. Maintenance of Transfection Rates and Physical Characterization of Lipid/DNA Complexes after Freeze-Drying and Rehydration. Archives of Biochemistry and Biophysics 348, 199–206 (1997).

37. Allison, S. D., Molina, M. d. C. & Anchordoquy, T. J. Stabilization of lipid/DNA complexes during the freezing step of the lyophilization process: the particle isolation hypothesis. Biochimica et Biophysica Acta (BBA) - Biomembranes 1468, 127–138 (2000).

38. Yadava, P., Gibbs, M., Castro, C. & Hughes, J. A. Effect of Lyophilization and Freeze-thawing on the Stability of siRNA-liposome Complexes. AAPS PharmSciTech 9, 335–341 (2007).

39. Grass, R. N., Heckel, R., Puddu, M., Paunescu, D. & Stark, W. J. Robust Chemical Preservation of Digital Information on DNA in Silica with Error-Correcting Codes. Angewandte Chemie International Edition 54, 2552–2555 (2015).

40. Erlich, Y. & Zielinski, D. DNA Fountain enables a robust and efficient storage architecture. Science 355, 950– 954 (2017).

41. Volkel, K. D. et al. FrameD: framework for DNA-based data storage design, verification, and validation. Bioinformatics 39, btad572 (2023).

42. Blawat, M. et al. Forward Error Correction for DNA Data Storage. Procedia Computer Science 80, 1011–1022 (2016).

43. Tabatabaei Yazdi, S. M. H., Yuan, Y., Ma, J., Zhao, H. & Milenkovic, O. A Rewritable, Random-Access DNA-Based Storage System. Sci Rep 5, 14138 (2015).

44. Chen, H. et al. Ionic strength-dependent persistence lengths of single-stranded RNA and DNA. Proceedings of the National Academy of Sciences 109, 799–804 (2012).

45. Stellwagen, E. & Stellwagen, N. C. Electrophoretic Mobility of DNA in Solutions of High Ionic Strength. Biophysical Journal 118, 2783–2789 (2020).

46. Oosawa, Fumio. Polyelectrolytes. (Marcel Dekker Inc., New York, NY, USA, 1971).

47. Ha, B.-Y. & Liu, A. J. Counterion-mediated, non-pairwise-additive attractions in bundles of like-charged rods. *Phys*. Rev. E 60, 803–813 (1999).

48. Lee, A. A., Perez-Martinez, C. S., Smith, A. M. & Perkin, S. Scaling Analysis of the Screening Length in Concentrated Electrolytes. Phys. Rev. Lett. 119, 026002 (2017).

49. Linse, P. & Lobaskin, V. Electrostatic Attraction and Phase Separation in Solutions of Like-Charged Colloidal Particles. Phys. Rev. Lett. 83, 4208–4211 (1999).

50. Kornyshev, A. A. & Leikin, S. Helical Structure Determines Different Susceptibilities of dsDNA, dsRNA, and tsDNA to Counterion-Induced Condensation. Biophysical Journal 104, 2031–2041 (2013).

51. Kornyshev, A. A. & Leikin, S. Electrostatic Zipper Motif for DNA Aggregation. Phys. Rev. Lett. 82, 4138– 4141 (1999).

52. Wu, Y.-Y., Zhang, Z.-L., Zhang, J.-S., Zhu, X.-L. & Tan, Z.-J. Multivalent ion-mediated nucleic acid helix-helix interactions: RNA versus DNA. Nucleic Acids Research 43, 6156–6165 (2015).

53. Kewalramani, S. et al. Electrolyte-Mediated Assembly of Charged Nanoparticles. ACS Cent. Sci. 2, 219–224 (2016).

54. Macfarlane, R. J. et al. Importance of the DNA “bond” in programmable nanoparticle crystallization. Proceedings of the National Academy of Sciences 111, 14995–15000 (2014).

55. Wang, M. X. et al. Altering DNA-Programmable Colloidal Crystallization Paths by Modulating Particle Repulsion. Nano Lett. 17, 5126–5132 (2017).

56. Bao, L., Zhang, X., Jin, L. & Tan, Z.-J. Flexibility of nucleic acids: From DNA to RNA*. Chinese Phys. B 25, 018703 (2015).

57. Ambia-Garrido, J., Vainrub, A. & Pettitt, B. M. A model for Structure and Thermodynamics of ssDNA and dsDNA Near a Surface: a Coarse Grained Approach. Comput Phys Commun 181, 2001–2007 (2010).

58. Singh, A., Maity, A. & Singh, N. Structure and Dynamics of dsDNA in Cell-like Environments. Entropy 24, 1587 (2022).

59. Ferreira, I., Amarante, T. D. & Weber, G. DNA terminal base pairs have weaker hydrogen bonds especially for AT under low salt concentration. The Journal of Chemical Physics 143, 175101 (2015).

60. Singh, A. & Singh, N. Effect of salt concentration on the stability of heterogeneous DNA. Physica A: Statistical Mechanics and its Applications 419, 328–334 (2015).

61. Tan, Z.-J. & Chen, S.-J. Salt Contribution to RNA Tertiary Structure Folding Stability. Biophys J 101, 176–187 (2011).

62. Sarkar, S., Maity, A., Sarma Phukon, A., Ghosh, S. & Chakrabarti, R. Salt Induced Structural Collapse, Swelling, and Signature of Aggregation of Two ssDNA Strands: Insights from Molecular Dynamics Simulation. J. Phys. Chem. B 123, 47–56 (2019).

63. Arscott, P. G., Li, A.-Z. & Bloomfield, V. A. Condensation of DNA by trivalent cations. 1. Effects of DNA length and topology on the size and shape of condensed particles. Biopolymers 30, 619–630 (1990).

64. Wong, G. C. L. & Pollack, L. Electrostatics of strongly charged biological polymers: ion-mediated interactions and self-organization in nucleic acids and proteins. Annu Rev Phys Chem 61, 171–189 (2010).

65. Korjus, K., Hebart, M. N. & Vicente, R. An Efficient Data Partitioning to Improve Classification Performance While Keeping Parameters Interpretable. PLOS ONE 11, e0161788 (2016).

66. Neagu, M.-I. & Miclea, L. Data scrambling in memories: A security measure. in 2014 *IEEE International Conference on Automation, Quality and Testing*, Robotics 1–6 (2014). doi:10.1109/AQTR.2014.6857847.

67. Benini, L., Galati, A., Macii, A., Macii, E. & Poncino, M. Energy-efficient data scrambling on memory-processor interfaces. in Proceedings of the 2003 international symposium on Low power electronics and design 26–29 (Association for Computing Machinery, New York, NY, USA, 2003). doi:10.1145/871506.871517.

