## Supplementary Information for "Colloidal DNA nanoaggregates applied towards file-partitioning for information storage and dynamic data obfuscation"

---

### Table of Contents

**Supplementary Notes**

Supplementary Note 1 | Important formulas used for plots in Fig. 5 & 6

**Supplementary Figures**

Supplementary Figure 1 | Formation of homogeneous aggregates of ssDNA and dsDNA

Supplementary Figure 2 | Reproducible homogeneous DNA nanoaggregates with 1M NaCl

Supplementary Figure 3 | Formation of homogeneous nanoaggregates with longer strands of DNA.

Supplementary Figure 4 | Formation of heterogeneous aggregates.

Supplementary Figure 5 | Size distributions of RNA nanoaggregates.

Supplementary Figure 6 | | Reproducible homogeneous DNA nanoaggregates with 1M $CaCl_2$.

Supplementary Figure 7 | Baseline DNA Bio-TEM images.

Supplementary Figure 8 | Additional BioTEM images for "nanobundles" and "nanoclusters".

Supplementary Figure 9 | Nanoaggregates increase in size with higher ionic strengths and in presence of divalent salts.

Supplementary Figure 10 | Thermally tunable aggregation system.

Supplementary Figure 11 | Schematic illustration of the magnetic separation process.

Supplementary Figure 12 | Characterization of nanoaggregates formed using file-encoded DNA + ovDNA.

Supplementary Figure 13 | Binary percentage of file-encoded DNA for each file in the mixture with ratio of pullout-oligo.

Supplementary Fig. 14 | Characterization of nanoaggregates formed using file-encoded DNA + ssDNA and ovDNA by changing PulloutDNA: File DNA ratios.

**Supplementary Tables**

Supplementary Table 1 | List of sequences used for experiments.

Supplementary Table 2 | Annotation of experimental conditions for Figure 6C and Figure 6D

Supplementary Table 3 | Annotation of experimental conditions for Figure 6F

Supplementary Table 4 | Annotation of experimental conditions for Supplementary Figure 19

Supplementary Table 5 | Annotation of experimental conditions for Figure 7a

Supplementary Table 6 | Annotation of experimental conditions for Figure 7b

#### Supplementary Note 1 | Formulas used for plots in Fig. 5 & 6

##### Percentage of Recovered DNA

In Figure 5, the percentage of recovered DNA was calculated by the following equation:

$$Percentage\ of\ recovered\ DNA = \frac{Amount\ of\ DNA\ measured\ in\ a\ solution}{Original\ amount\ of\ the\ same\ DNA}$$

##### File recovery percentage

In Figure 6, the file recovery percentage was calculated by the following equation:

$$File\ recovery\ percentage = \frac{Amount\ of\ the\ selected\ files\ strads\ \ in\ a\ solution}{Original\ amount\ of\ the\ same\ file\ strands}$$

##### Percentage of file in the mixture

In Figure 6, the percentage of file in the mixture was calculated by the following equation:

$$Percentage\ of\ file\ in\ the\ mixture = \frac{\begin{array}{c}Amount\ of\ the\ selected\ file\ strands\\ in\ a\ solution\end{array}}{\begin{array}{c}Total\ amount\ of\ file\ strands\\ in\ the\ same\ solution\end{array}}$$

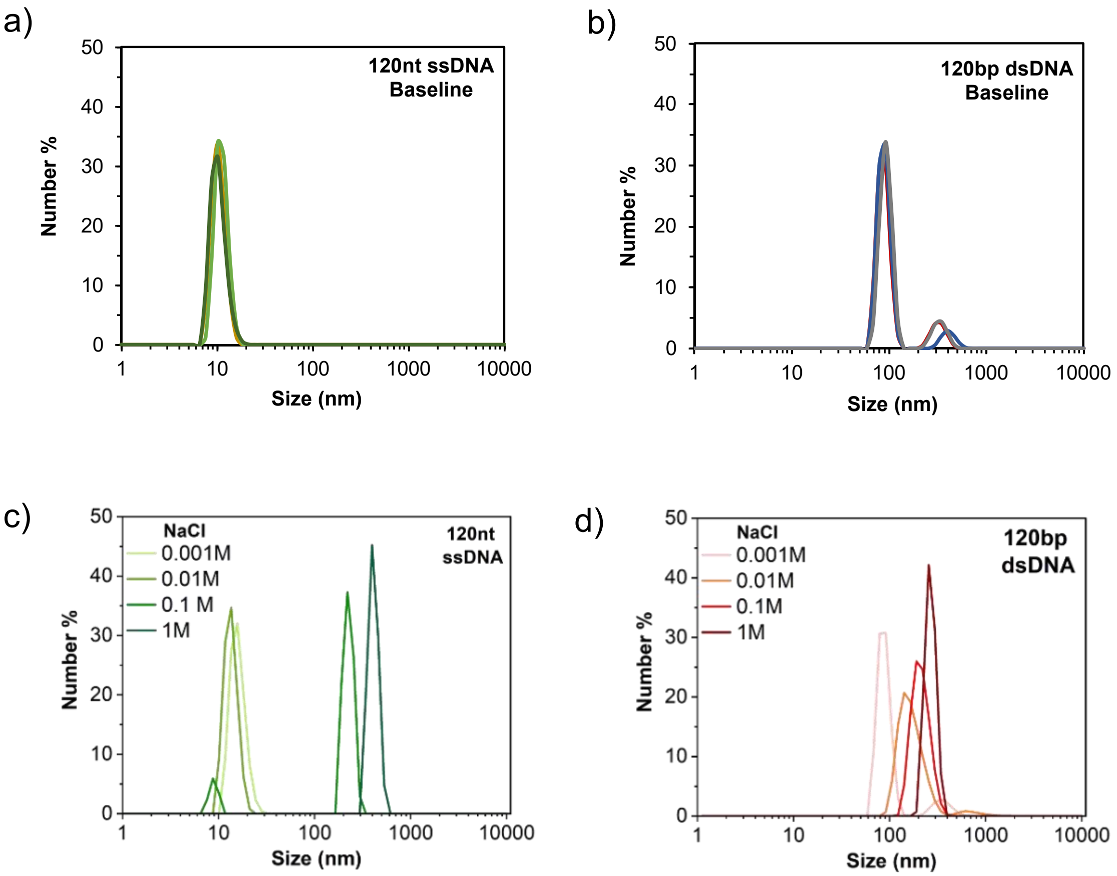


**Supplementary Fig. 1 | Formation of homogeneous aggregates of ssDNA and dsDNA.** Plots show average baseline measurements of DNA of sizes **a,** 120nt ssDNA **b,** 120bp ds DNAs. After these measurements, DNAs are subjected to solutions with increasing concentration of NaCl. Salt actuated aggregation starts as the concentration moves towards the higher range. Plots with **c,** 120nt ssDNA and **d,** 120bp dsDNA show the shifting of peaks in light scattering.

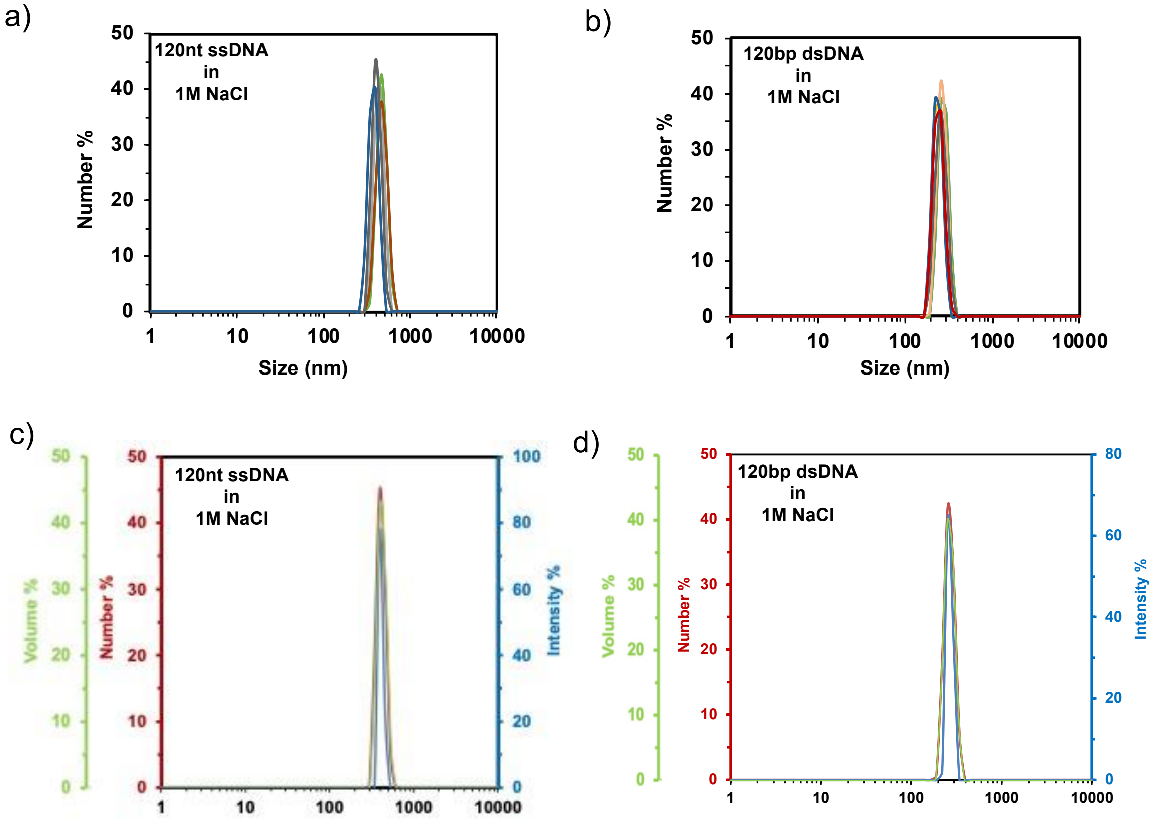


**Supplementary Fig. 2 | Formation of reproducible homogeneous DNA nanoaggregates in 1M NaCl**. **a,** 120ss **b,** 120ds The peaks overlap indicating the formation of nanoaggregates is consistent and reproducible. Plots with average measurements are included in the main manuscript.

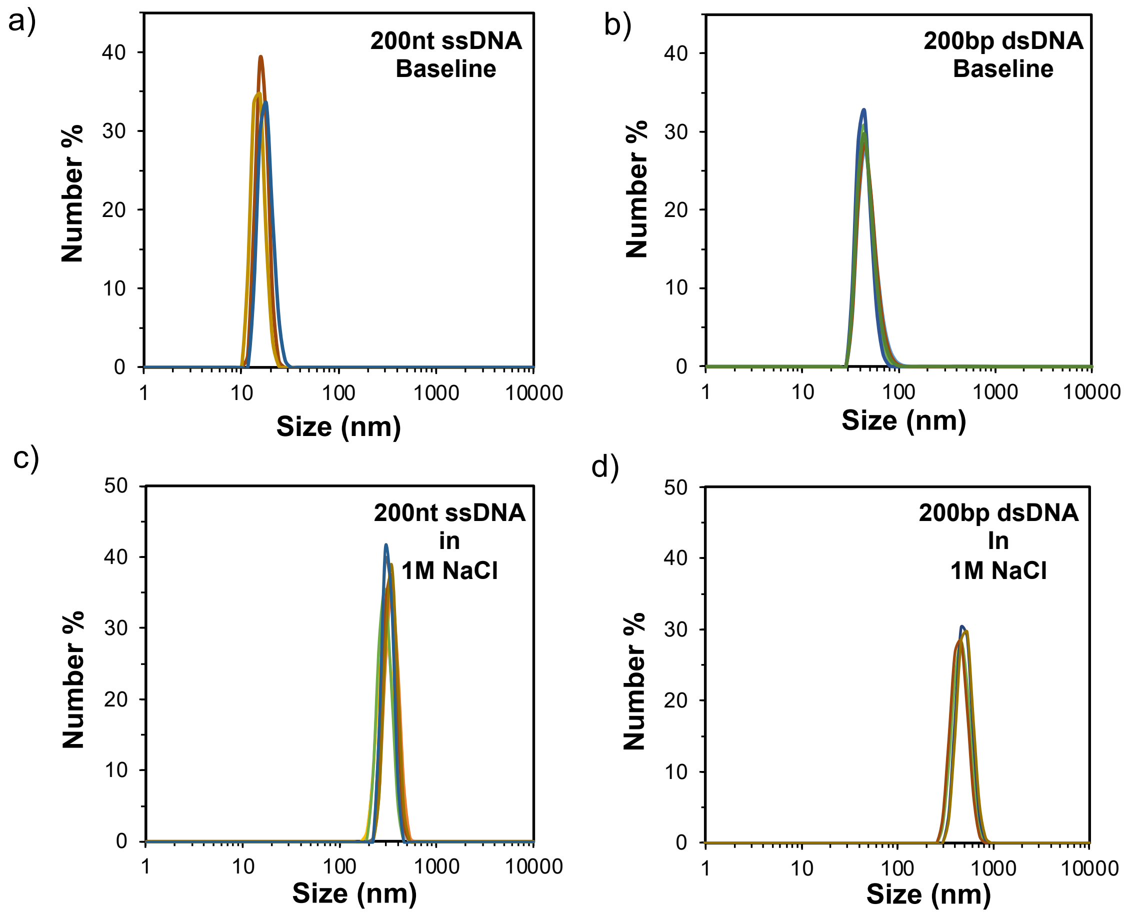


**Supplementary Fig. 3 | Formation of homogeneous nanoaggregates with longer strands of DNA.** Plots show average baseline measurements of DNA of sizes **a,** 200nt ss **b,** 200bp ds DNAs. After these measurements, **c,** 200nt ss **d,** 200bp ds DNAs were subjected to 1M NaCl. Consistent results observed for multiple samples.

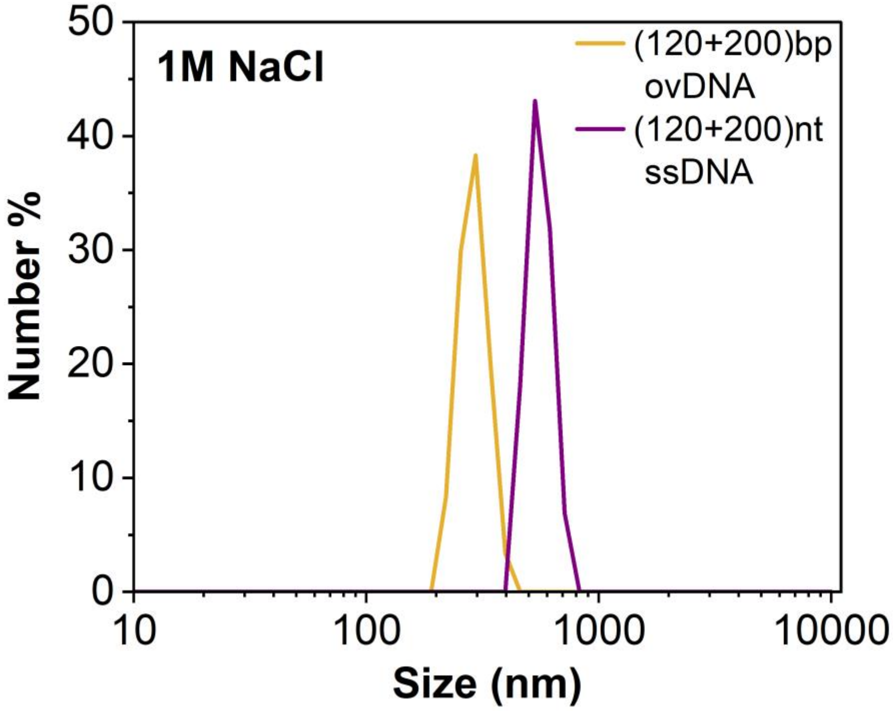


**Supplementary Fig. 4 | Formation of heterogeneous aggregates.** These aggregates are formed by initially mixing two different lengths of DNA and then subjecting the mixture to a high salt environment to initiate aggregation.

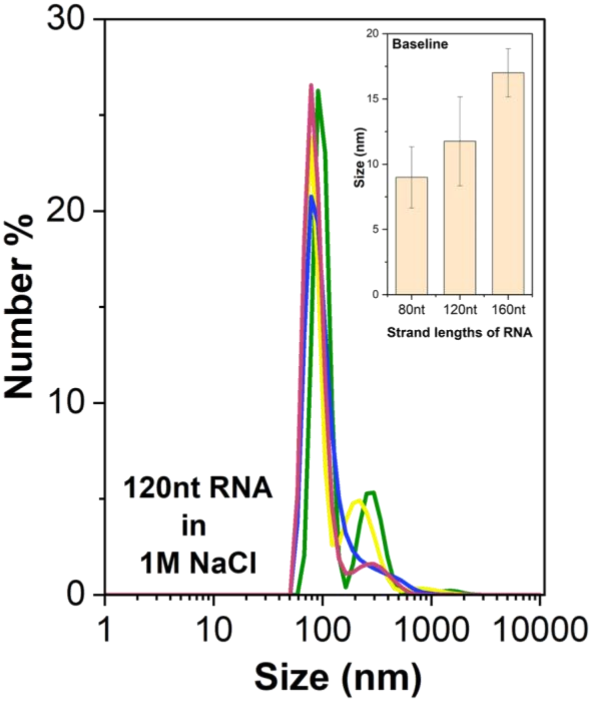


**Supplementary Fig. 5 | Size distributions of RNA nanoaggregates.** Multiple measurements of RNA aggregates indicate some fluctuations, revealing that RNA nanoaggregates are less consistent and stable compared to DNA aggregates.

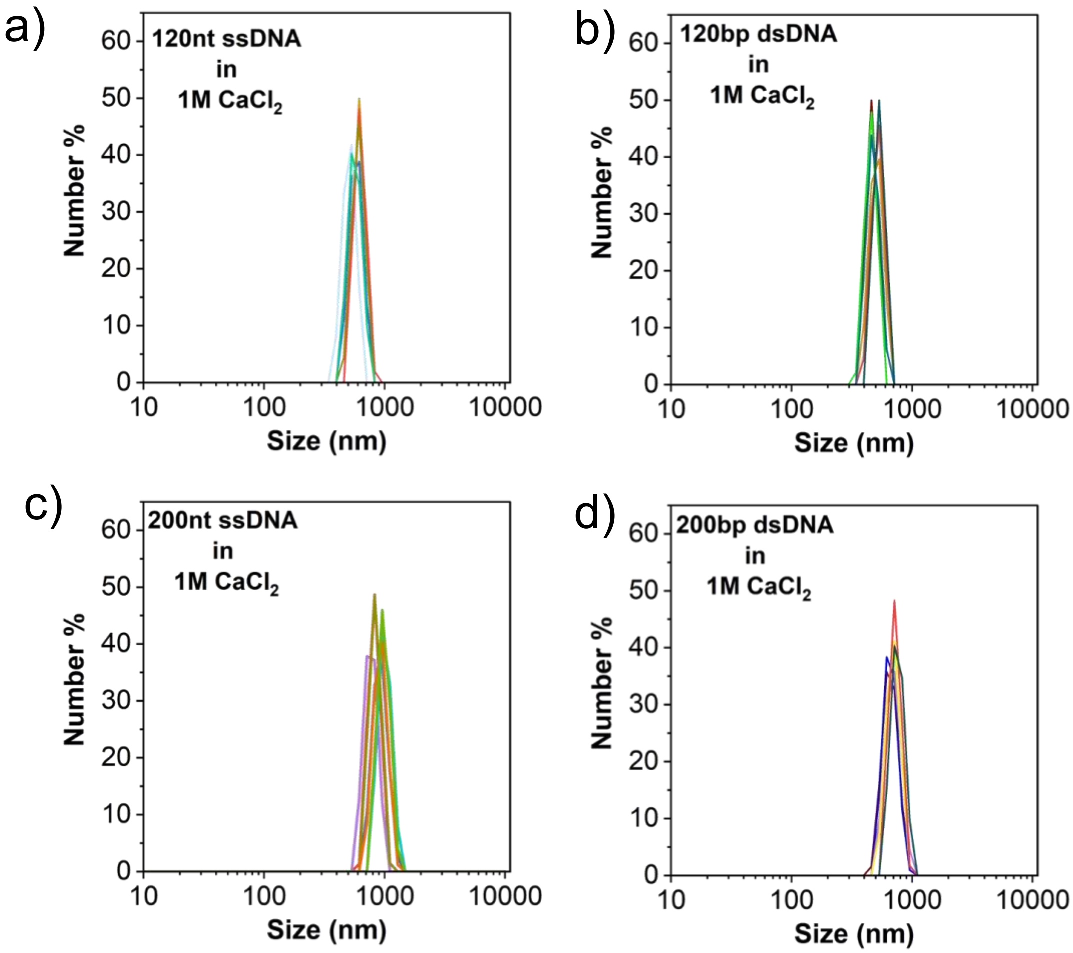


**Supplementary Fig. 6 | Reproducible homogeneous DNA nanoaggregation in 1M $CaCl_2$. a,** 120ss **b,** 120ds **c**, 200ss and **d,** 200ds. The peaks overlap indicating the formation of nanoaggregates is consistent and reproducible. Plots with average measurements are included in the main manuscript.

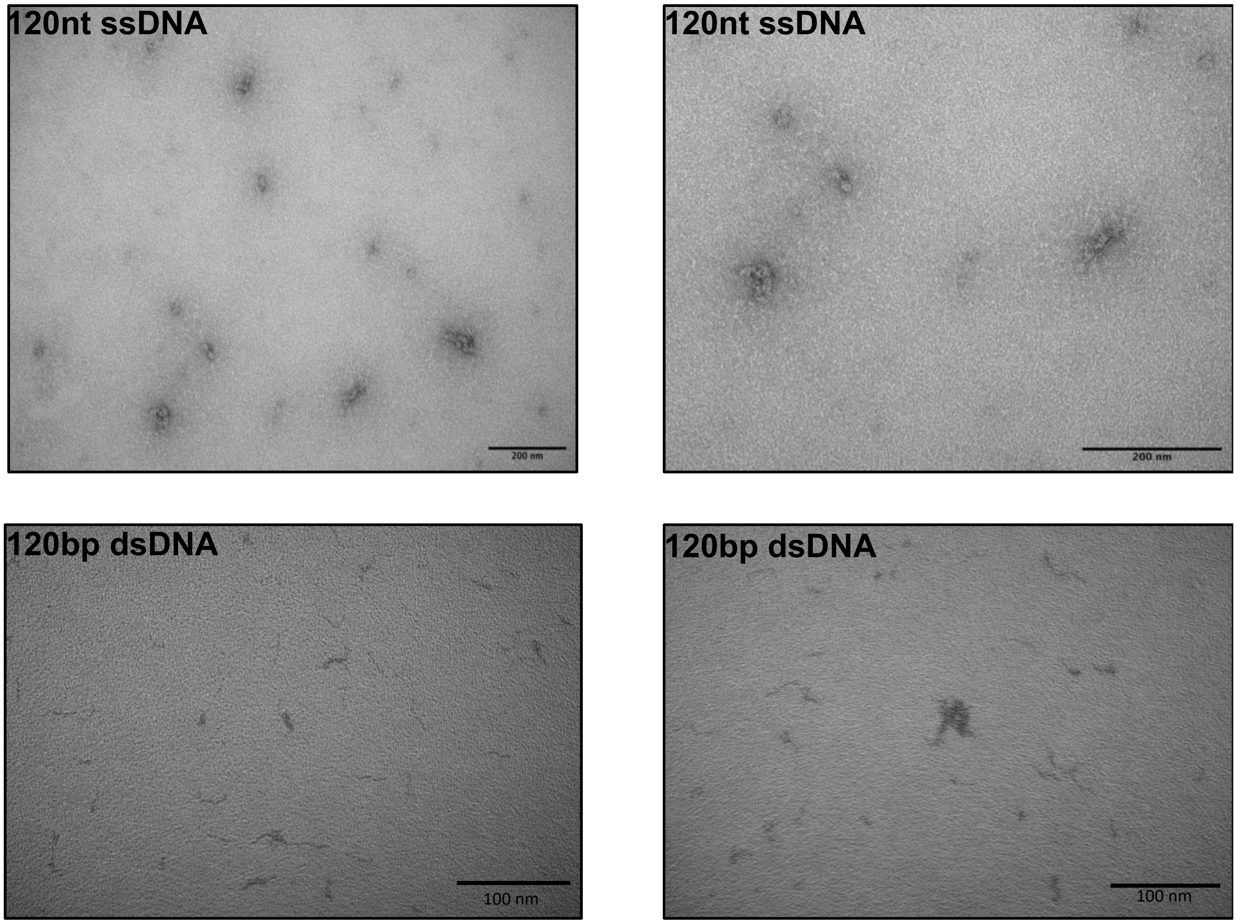


**Supplementary Fig. 7 | Baseline DNA Bio-TEM images** - 120nt ssDNA (top row) and 120bp dsDNA (bottom row). Scalebars are 200 nm for top row and 100 nm for bottom row.

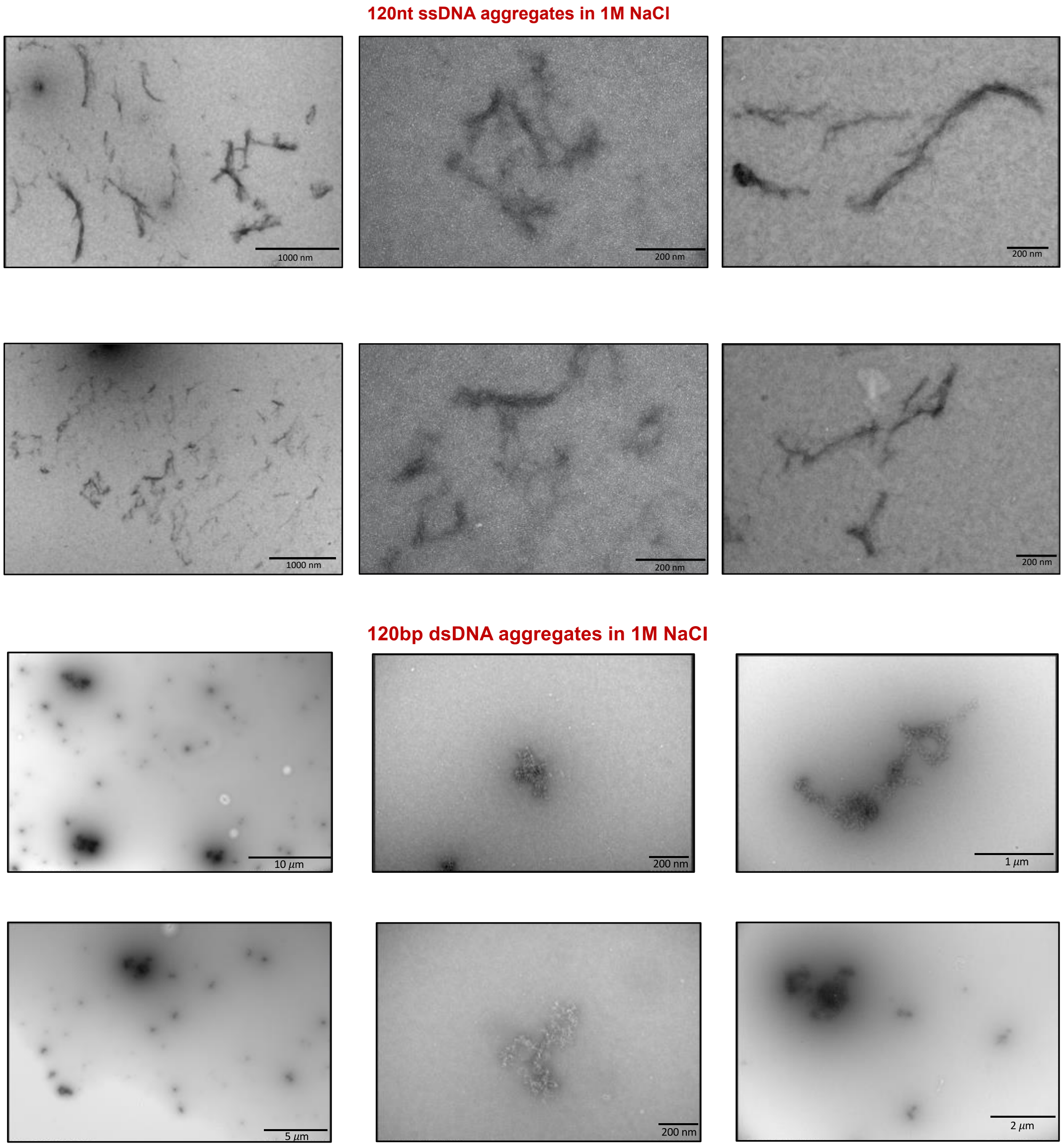


**Supplementary Fig. 8 | Additional BioTEM images for "nanobundles" and "nanoclusters".** The first column displays images at lower magnification, indicating that the aggregates are uniformly distributed throughout the area. Some aggregates appear more clustered than others, which could be attributed to staining defects introduced during sample preparation. Certain sections reveal that multiple aggregates overlap or are interconnected.

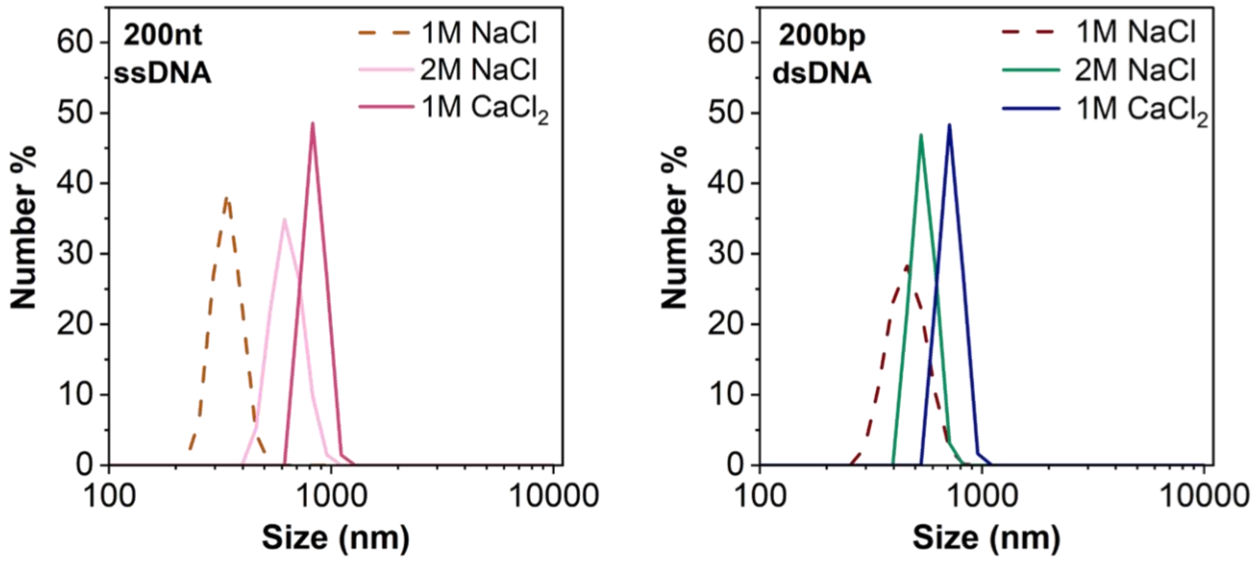


**Supplementary Fig. 9 | Nanoaggregates increase in size with higher ionic strengths and in presence of divalent salts.** Plots show the size comparison of aggregates formed in the presence of 1M NaCl, 2M NaCl and 1M $CaCl_2$ for ss and dsDNA with a higher strand length of 200.

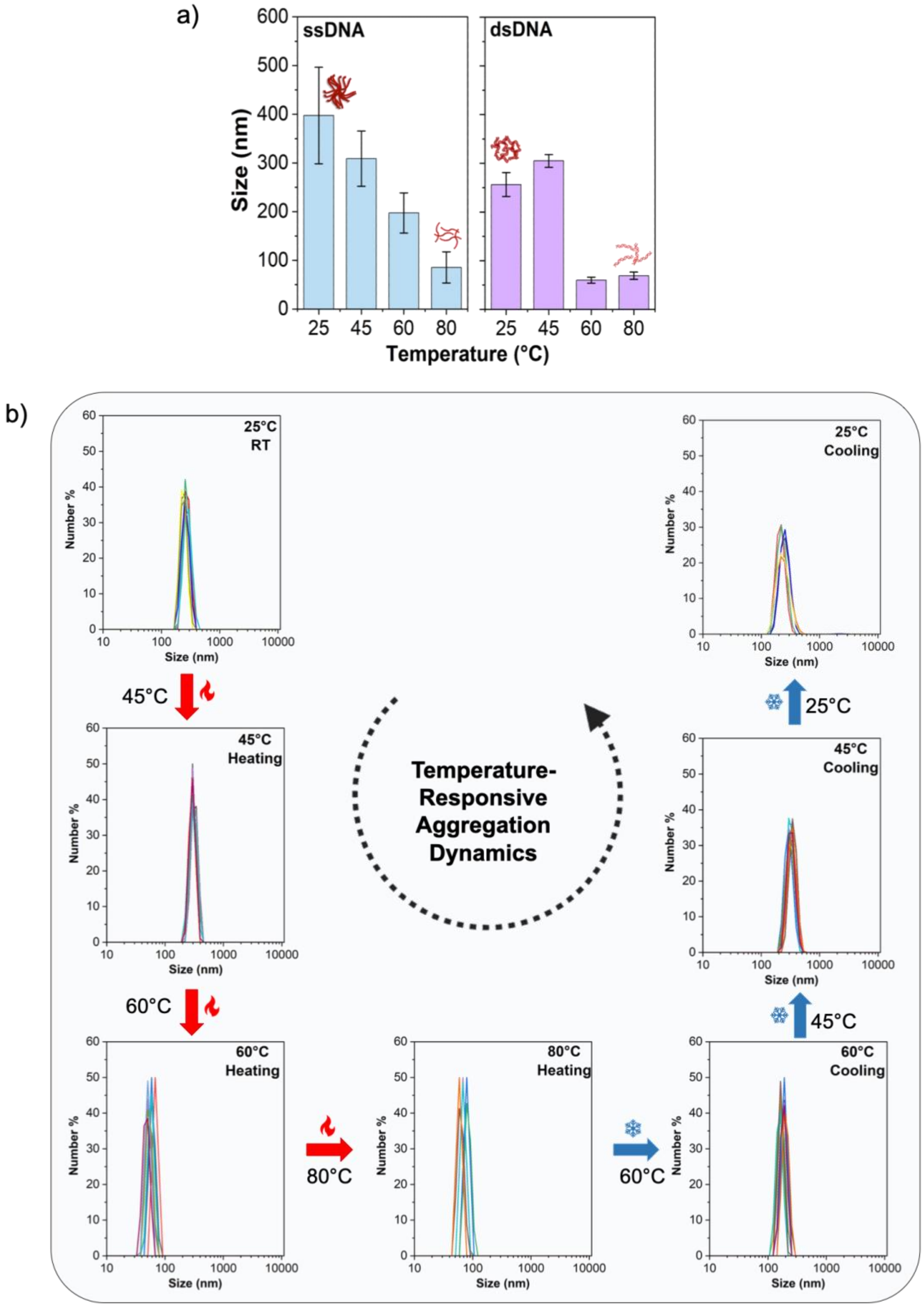


**Supplementary Fig. 10| Thermally tunable aggregation system. a,** Temperature-responsive aggregation dynamics for ss- and dsDNA. Aggregates are stable till 45°C and are observed to break apart at 60°C and remain in that state through 80°C. **b,** Reversible and dynamic aggregation with 120 bp dsDNA. This study was conducted in a continuous setting. Initially, the sample is measured at ambient temperature. Then, the temperature of the system was increased to 45°C, maintaining consistent conditions. Multiple measurements were taken to ensure repeatability. Following the measurements at 45°C, the temperature was further raised to 60°C and then to 80°C respectively. The system was allowed to equilibrate at each temperature before starting any measurements. After measuring at 80°C, the temperature was gradually lowered to 60°C, then to 45°C and finally back to 25°C in a reverse cycle. This procedure is designed to observe the changes in nature of the aggregates during the process of heating and cooling. Aggregates are observed to break apart at 60°C and remain in that state through 80°C. Upon cooling, the aggregates reformed, and after returning to room temperature, we observe aggregates of similar size to those initially present.

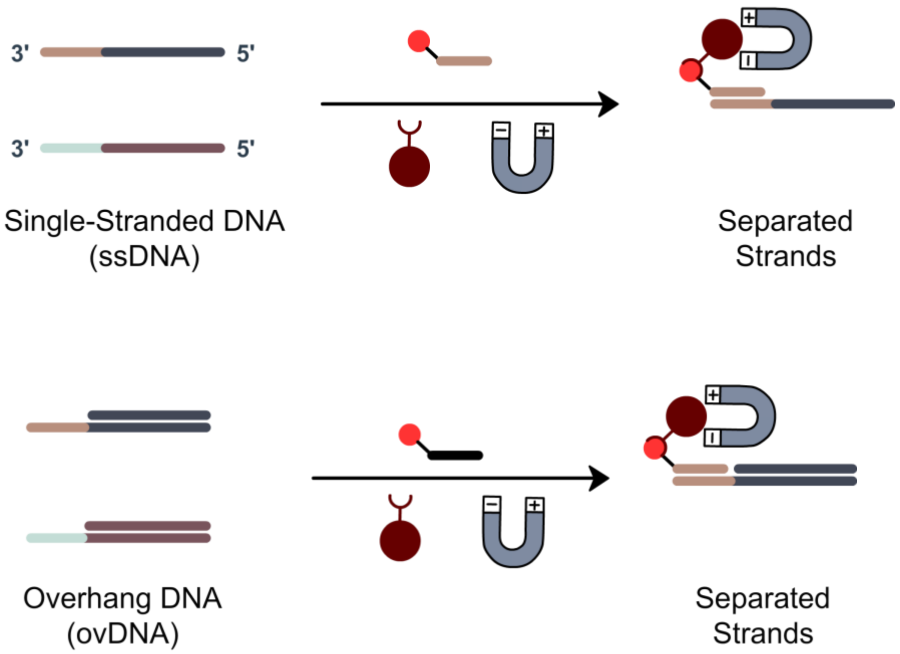


**Supplementary Fig. 11 | Schematic illustration of the magnetic separation process. ssDNA** (top row) and **ovDNA** (bottom row). This separation method was used to assess the strand integrity of the DNA nanoaggregates.

a)

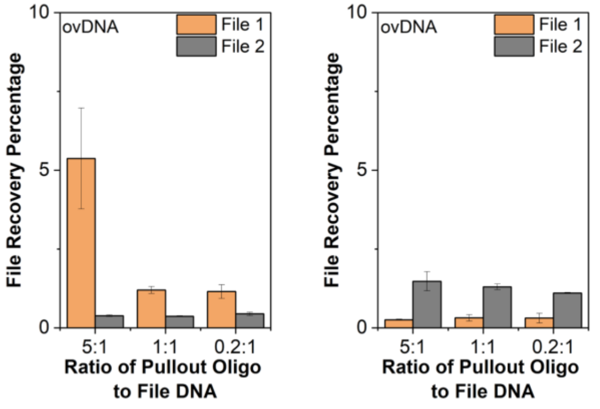


| PulloutDNA:FileDNA | Operation (Left) | Operation (Right) |
|---|---|---|
| 5:1 | Agg(ovDNA+File1) + Agg (File2) + Non-Vortexing | Agg(ovDNA+File2) + Agg (File1) + Non- Vortexing |
| 1:1 | Agg(ovDNA+File1) + Agg (File2) + Non-Vortexing | Agg(ovDNA+File2) + Agg (File1) + Non- Vortexing |
| 0.2:1 | Agg(ovDNA+File1) + Agg (File2) + Non-Vortexing | Agg(ovDNA+File2) + Agg (File1) + Non- Vortexing |

b)

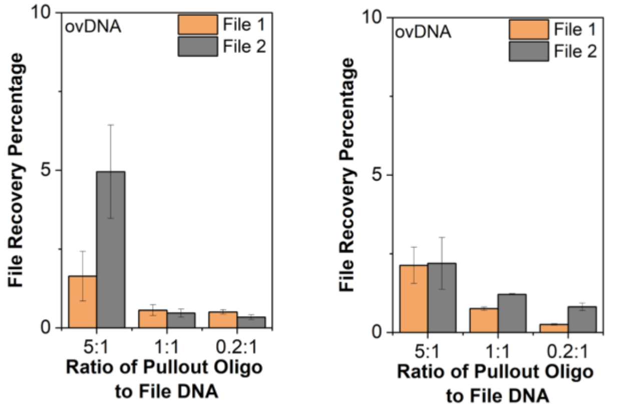


| PulloutDNA:FileDNA | Operation (Left) | Operation (Right) |
|---|---|---|
| 5:1 | Agg(ovDNA+File1) + Agg (File2) + Vortexing | Agg(ovDNA+File2) + Agg (File1) + Vortexing |
| 1:1 | Agg(ovDNA+File1) + Agg (File2) + Vortexing | Agg(ovDNA+File2) + Agg (File1) + Vortexing |
| 0.2:1 | Agg(ovDNA+File1) + Agg (File2) + Vortexing | Agg(ovDNA+File2) + Agg (File1) + Vortexing |

**Supplementary Fig. 12| Characterization of nanoaggregates formed using file-encoded DNA + ovDNA.** Quantification of DNA for each file-encoded DNA in elution with varying ratios of pullout-oligo: file-encoded-DNA, ranging from 5:1, 1:1 and 0.2:1 (see corresponding Tables for annotation of experimental conditions) in two cases. **a,** non-vortex and **b,** with vortex. The ratio of 5:1 yielded the best recovery.

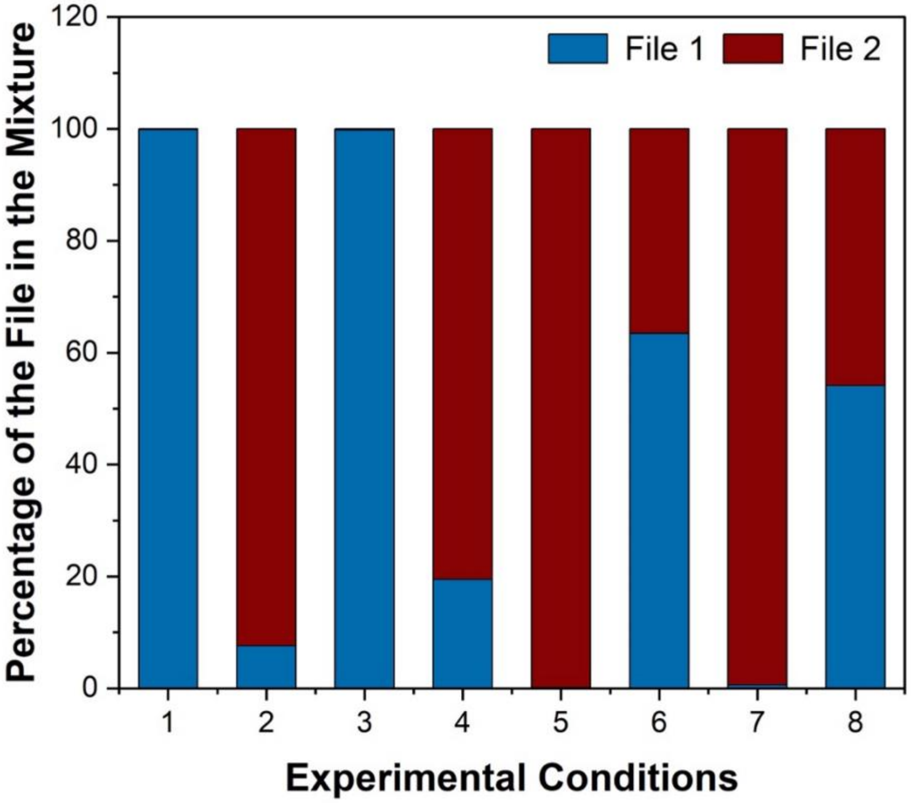


**Supplementary Fig. 13 | Binary percentage of file-encoded DNA for each file in the mixture with ratio of pullout-oligo: file-encoded-DNA at 5:1.** See corresponding Table S4 for annotation of experimental conditions.

a)

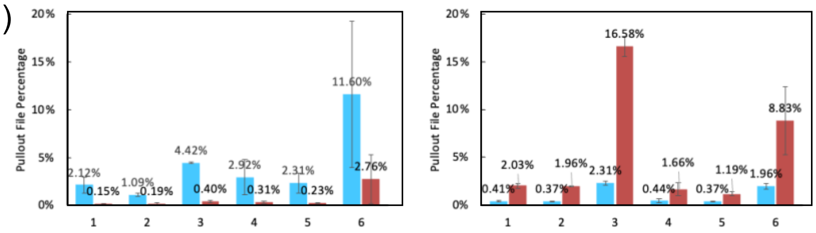


| ID | PulloutDNA:File DNA | Operation (Left) | Operation (Right) |
|---|---|---|---|
| 1 | 1:0.2 | Agg(ssDNA+File1) + Agg (File2) + Non-Vortexing | Agg(ssDNA+File2) + Agg (File1) + Non-Vortexing |
| 2 | 1:1 | Agg(ssDNA+File1) + Agg (File2) + Non-Vortexing | Agg(ssDNA+File2) + Agg (File1) + Non-Vortexing |
| 3 | 1:5 | Agg(ssDNA+File1) + Agg (File2) + Non-Vortexing | Agg(ssDNA+File2) + Agg (File1) + Non-Vortexing |
| 4 | 1:0.2 | Agg(ovDNA+File1) + Agg (File2) + Non-Vortexing | Agg(ovDNA+File2) + Agg (File1) +Non-Vortexing |
| 5 | 1:1 | Agg(ovDNA+File1) + Agg (File2) + Non-Vortexing | Agg(ovDNA+File2) + Agg (File1) + Non-Vortexing |
| 6 | 1:5 | Agg(ovDNA+File1) + Agg (File2) + Non-Vortexing | Agg(ovDNA+File2) + Agg (File1) + Non-Vortexing |

b)

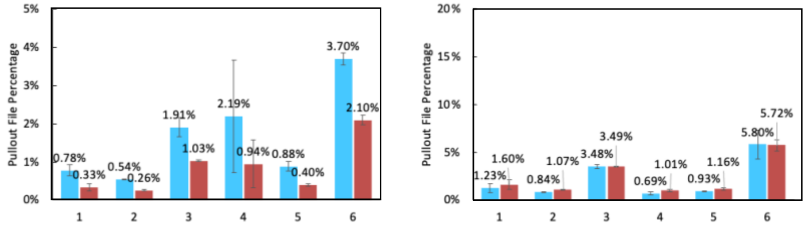


| ID | PulloutDNA:File DNA | Operation (Left) | Operation (Right) |
|---|---|---|---|
| 1 | 1:0.2 | Agg(ssDNA+File1) + Agg (File2) + Vortexing | Agg(ssDNA+File2) + Agg (File1) + Vortexing |
| 2 | 1:1 | Agg(ssDNA+File1) + Agg (File2) + Vortexing | Agg(ssDNA+File2) + Agg (File1) + Vortexing |
| 3 | 1:5 | Agg(ssDNA+File1) + Agg (File2) + Vortexing | Agg(ssDNA+File2) + Agg (File1) + Vortexing |
| 4 | 1:0.2 | Agg(ovDNA+File1) + Agg (File2) + Vortexing | Agg(ovDNA+File2) + Agg (File1) + Vortexing |
| 5 | 1:1 | Agg(ovDNA+File1) + Agg (File2) + Vortexing | Agg(ovDNA+File2) + Agg (File1) + Vortexing |
| 6 | 1:5 | Agg(ovDNA+File1) + Agg (File2) + Vortexing | Agg(ovDNA+File2) + Agg (File1) + Vortexing |

**Supplementary Fig. 14 | Characterization of nanoaggregates formed using file-encoded DNA + ssDNA and ovDNA by changing PulloutDNA: File DNA ratios.** Quantification of DNA for each file-encoded DNA in elution with varying ratios of pullout-oligo: file-encoded-DNA, ranging from 1:0.2, 1:1 and 1:5 (see corresponding Tables for annotation of experimental conditions) in two cases. **a,** non-vortex and **b,** with vortex.

#### Supplementary Table 1 | List of sequences used for experiments.

| ssDNA samples | |
|---|---|
| **Sample** | **Sequence** |
| 120 ssDNA | ggatagaaggtggagccgaaggatgtttttgatagcatagcgttgctgtcagacctaacctaccatgactagaagctccagcacaacgttcttccggatacccgcagcgaaaatcacgtc |
| 200 ssDNA | ggatagaaggtggagccgaaggatgtttttgatagcatagcgttgctgtcagacctaacctaccatgactagaagctccagcacaacgttcttccggatacccgcagcgaaaatcacgtcggatagaaggtggagccgaaggatgtttttgatagcatagcgttgctgtcagacctaacctaccatgactagaagctcca |
| 120 ssDNA 0% GC | atatatatattataaaattataaaaaaaaaaaaaaataataaattataaattattattaatttataaataatttaataaataattttaaatataaatattaaaaaaaattttaattatta |
| 120 ssDNA 25% GC | gatttttattaacgcttaatctatcatagttattcccatatttaaatttgaacataggttttgtcagggataattctaatcactaactaaattatctattaataagaatctataatattt |
| 120 ssDNA 50% GC | cagagtcatcaccgtcttctcctccgtctccacgcagagtgtcgattaccagcttcttaagcgagccttaagcctaaattgaattcaccaggcccatacgcgcagcactcgtaaatgtta |
| 120 ssDNA 75% GC | cggcttacacgcgtgttagggcgagctgagaccgcgggtccgccgggcccccgggccgcctcggccatccgcgcaccctcggcactcccgctccgaccgcggagaaggtaccggtccccg |

| dsDNA samples | |
|---|---|
| **Sample** | **Sequence** |
| 120 dsDNA | gggagtaatccccttggcggtcgcgggggacagcgcgtacgtgcgtttaagcggtgctagagctgtctacgaccagcgcgcgctatagtgagtcgtattaggattctccagggcatccgg |
| Oligos needed to make dsDNA: | ccggatgccctggagaatcc |
| | gggagtaatccccttggcggt |
| Oligo needed to make ovDNA: | taatacgactcactatagcgcgc |

| Sample | Sequence |
|---|---|
| 200 dsDNA | caggtacgcagttagcactccgtacgtacgtacgcagctagctcgatgagtactctgctcgatgagtactctgctcgacgagatgagacgagtctctcgtagacgagagcagactcagtcatcgcgctagagagcatagagtcgtgatctatgctcagcgcgctatagtgagtcgtattatccgtagtcatattgccacg |
| Oligos needed to make dsDNA: | cgtggcaatatgactacgga |
| | caggtacgcagttagcactc |
| Oligo needed to make ovDNA | taatacgactcactatagcgcgc |

| RNA samples | |
|---|---|
| **Sample** | **DNA Sequence for making RNA** |
| 80nt RNA | caggtacgcagttagcactcagtctctcgtagacgagagcagactcagtcatcgcgctagagagcatagagtcgtgagcgcgctatagtgagtcgtattatccgtagtcatattgccacg |
| 120nt RNA | caggtacgcagttagcactctactctgctcgatgagtactctgctcgacgagatgagacgagtctctcgtagacgagagcagactcagtcatcgcgctagagagcatagagtcgtgagcgcgctatagtgagtcgtattatccgtagtcatattgccacg |
| 160nt RNA | caggtacgcagttagcactccgtacgtacgtacgcagctagctcgatgagtactctgctcgatgagtactctgctcgacgagatgagacgagtctctcgtagacgagagcagactcagtcatcgcgctagagagcatagagtcgtgatctatgctcagcgcgctatagtgagtcgtattatccgtagtcatattgccacg |

**Supplementary Table 2 | Annotation of experimental conditions for Figure 6C and Figure 6D**

| ID | PulloutDNA: FileDNA | Operation (6C) | Operation (6D) |
|---|---|---|---|
| 1 | 5:1 | Agg(ssDNA-S1+File1) + Agg (File2) + Non-Vortexing | Agg(ssDNA-S2+File2) + Agg (File1) +Non- Vortexing |
| 2 | 1:1 | Agg(ssDNA-S1+File1) + Agg (File2) + Non-Vortexing | Agg(ssDNA-S2+File2) + Agg (File1) + Non- Vortexing |
| 3 | 0.2:1 | Agg(ssDNA-S1+File1) + Agg (File2) + Non-Vortexing | Agg(ssDNA-S2+File2) + Agg (File1) + Non- Vortexing |

**Supplementary Table 3 | Annotation of experimental conditions for Figure 6F**

| ID | PulloutDNA: FileDNA | Operation (6F Left) | Operation (6F- Right) |
|---|---|---|---|
| 1 | 5:1 | Agg(ssDNA-S1+File1) + Agg (File2) + Vortexing | Agg(ssDNA-S2+File2) + Agg (File1) + Vortexing |
| 2 | 1:1 | Agg(ssDNA-S1+File1) + Agg (File2) + Vortexing | Agg(ssDNA-S2+File2) + Agg (File1) + Vortexing |
| 3 | 0.2:1 | Agg(ssDNA-S1+File1) + Agg (File2) + Vortexing | Agg(ssDNA-S2+File2) + Agg (File1) + Vortexing |

**Supplementary Table 4 | Annotation of experimental conditions for Supplementary Figure 19**

| ID | PulloutDNA: FileDNA | Pullout Oligo | Operation |
|---|---|---|---|
| 1 | 5:1 | ssS1 | Agg(ssDNA-S1+File1) + Agg (File2) + Non-Vortexing |
| 2 | 5:1 | ssS1 | Agg(ssDNA-S1+File1) + Agg (File2) + Vortexing |
| 3 | 5:1 | ovS1 | Agg(ovDNA-S1+File1) + Agg (File2) + Non-Vortexing |
| 4 | 5:1 | ovS1 | Agg(ovDNA-S1+File1) + Agg (File2) + Vortexing |
| 5 | 5:1 | ssS2 | Agg(ssDNA-S2+File2) + Agg (File1) + Non-Vortexing |
| 6 | 5:1 | ssS2 | Agg(ssDNA-S2+File2) + Agg (File1) + Vortexing |
| 7 | 5:1 | ovS2 | Agg(ovDNA-S2+File2) + Agg (File1) + Non-Vortexing |
| 8 | 5:1 | ovS2 | Agg(ovDNA-S2+File2) + Agg (File1) + Vortexing |

**Supplementary Table 5 | Annotation of experimental conditions for Figure 7a**

| ID | Pullout oligo | Ratio of pullout-oligo to file DNA | Mixing Order | Vortex | Targeted File | Decoding File |
|---|---|---|---|---|---|---|
| 1 | Bio-ssS1 | 5:1 | Agg(bio-ssS1+File1) + Agg(file2) | No | File1 | File1 |
| 2 | Bio-ssS1 | 5:1 | Agg(bio-ssS1+File1) + Agg(file2) | Yes | File1 | File1 |
| 3 | Bio-ovS1 | 5:1 | Agg(bio-ovS1+File1) + Agg(file2) | No | File1 | File1 |
| 4 | Bio-ovS1 | 5:1 | Agg(bio-ovS1+File1) + Agg(file2) | Yes | File1 | File1 |
| 5 | Bio-ssS2 | 5:1 | Agg(bio-ssS2+File2) + Agg(file1) | No | File2 | File1 |
| 6 | Bio-ssS2 | 5:1 | Agg(bio-ssS2+File2) + Agg(file1) | Yes | File2 | File1 |
| 7 | Bio-ovS2 | 5:1 | Agg(bio-ovS2+File2) + Agg(file1) | No | File2 | File1 |
| 8 | Bio-ovS2 | 5:1 | Agg(bio-ovS2+File2) + Agg(file1) | Yes | File2 | File1 |

**Supplementary Table 6 | Annotation of experimental conditions for Figure 7b**

| ID | Pullout oligo | Ratio of pullout-oligo to file DNA | Mixing Order | Vortex | Targeted File | Decoding File |
|---|---|---|---|---|---|---|
| 1 | Bio-ssS1 | 5:1 | Agg(bio-ssS1+File1)<br>+<br>Agg(file2) | No | File1 | File2 |
| 2 | Bio-ssS1 | 5:1 | Agg(bio-ssS1+File1)<br>+<br>Agg(file2) | Yes | File1 | File2 |
| 3 | Bio-ovS1 | 5:1 | Agg(bio-ovS1+File1)<br>+<br>Agg(file2) | No | File1 | File2 |
| 4 | Bio-ovS1 | 5:1 | Agg(bio-ovS1+File1)<br>+<br>Agg(file2) | Yes | File1 | File2 |
| 5 | Bio-ssS2 | 5:1 | Agg(bio-ssS2+File2)<br>+<br>Agg(file1) | No | File2 | File2 |
| 6 | Bio-ssS2 | 5:1 | Agg(bio-ssS2+File2)<br>+<br>Agg(file1) | Yes | File2 | File2 |
| 7 | Bio-ovS2 | 5:1 | Agg(bio-ovS2+File2)<br>+<br>Agg(file1) | No | File2 | File2 |
| 8 | Bio-ovS2 | 5:1 | Agg(bio-ovS2+File2)<br>+<br>Agg(file1) | Yes | File2 | File2 |
